# Optogenetic SH3 nanoclustering of SRC and HCK uncovers the functional specificity of these redundant kinases in macrophages

**DOI:** 10.64898/2025.12.27.692121

**Authors:** Cristina Torres-Torres, Paul Rivier, Satish Babu Moparthi, Benjamin Graedel, Lucien Hinderling, Lucia Campos-Perello, Christiane Oddou, Ingrid Bourin-Reynard, Alexei Grichine, Eva Faurobert, Emmanuelle Planus, Fabrice Senger, Olivier Pertz, Stéphane Vassilopoulos, Olivier Destaing

**Author notes:** To whom correspondence should be addressed, **Institute for Advanced Biosciences, Centre de Recherche UGA / Inserm U 1209 / CNRS UMR 5309, Site Santé - Allée des Alpes 38700 La Tronche**, France. Equal contributions (Cristina Torres-Torres, Paul Rivier). Equal contributions (Benjamin Graedel, Lucien Hinderling).

## Abstract

Understanding redundancy among Src Family Kinases (SFKs) challenges how cells achieve signaling specificity using closely related enzymes, and paves the way to revisit a basic property of any cell signaling network. Optogenetic control of the adaptor properties of the most redundant SFK members, SRC and HCK, was used to examine their potential specificity in macrophages. Uncoupling their kinase and adaptor functions uncovers an unique mechanism driven by oligomerized- and SH3-dependent specific binding that enables inducing distinct signaling outputs for each kinase. OptoSRC and optoHCK probes revealed that SRC and HCK can trigger specific cellular responses, despite only supposed overlapping functions. The specificity of optoHCK even unveils a novel role in activating polarized clathrin hotspots essential for directional migration. Redundancy could be rebuilt within optoHCK through stepwise optogenetic reconstruction of adaptor modules revealing the essential role of both membrane anchoring and SH2-domain. This deconstructive approach of SFK activation elucidated the molecular basis of their shared and unique signaling activities and supports a model of cooperative functional entanglement between SFK members, rather than simple redundancy.

## Introduction

Redundancy is a central property of signaling networks that appears reinforced along evolution, complicating our understanding of the genesis of specific cellular responses. At the molecular level, redundancy arises from the co-expression of multiple members of conserved protein families implicated in signaling that can elicit similar cellular responses^1^. It warrants a high level of resilience of signaling networks likely by enhancing their efficiency in transmitting information through noisy environment^2,3^. As a drawback, redundancy poses a significant challenge for therapeutic targeting, since compensatory pathways can reduce the effectiveness of drugs used to treat chronic diseases and cancer. Understanding the molecular entanglement between redundancy and specificity -how individual protein family members make distinct signaling decision-making events-remains a challenge. This necessitates revisiting from dynamic, functional, and mechanistic perspectives how signaling networks encode information and coordinate complex cellular behaviors.

The SRC family kinases (SFKs) offer a particularly relevant model as they belong to one of the largest kinase families. SFKs consist of eight members with complex expression pattern across tissues^4^. Three members (SRC, FYN and YES) are ubiquitously expressed and five members (HCK, FYN, FGR, BLK, LCK) are primarily enriched in hematopoietic cells (Fig. 1A). Despite SRC being the first discovered proto-oncogene, genetic studies revealed a surprising resilience of mouse models upon its loss^5,6^. In the myeloid lineage, one of the most SRC-sensitive cell types, the loss of SRC is largely compensated by HCK, suggesting a strong degree of functional redundancy among these two kinases^7^. As SRC, HCK has been implicated in the regulation of cell morphology^4^, migration and invasion processes^8^. In particular, both kinases modulate podosomes, specialized actomyosin-rich adhesion structures that support the invasive behavior of myeloid cells^9–11^. Beyond physiology, HCK overexpression is strongly linked to increased invasiveness in several solid tumors and lymphomas^12,13^, similarly to SRC^14^. Given that macrophages co-express up to six distinct SFKs (SRC, FYN, YES, HCK, FGR and LYN), their signaling network offers a powerful model to interrogate whether apparent redundancy could mask finely tuned specificity, helping to revisit the complexity of functional redundancy and potential specialization in macrophage signaling.

**Figure 1:**
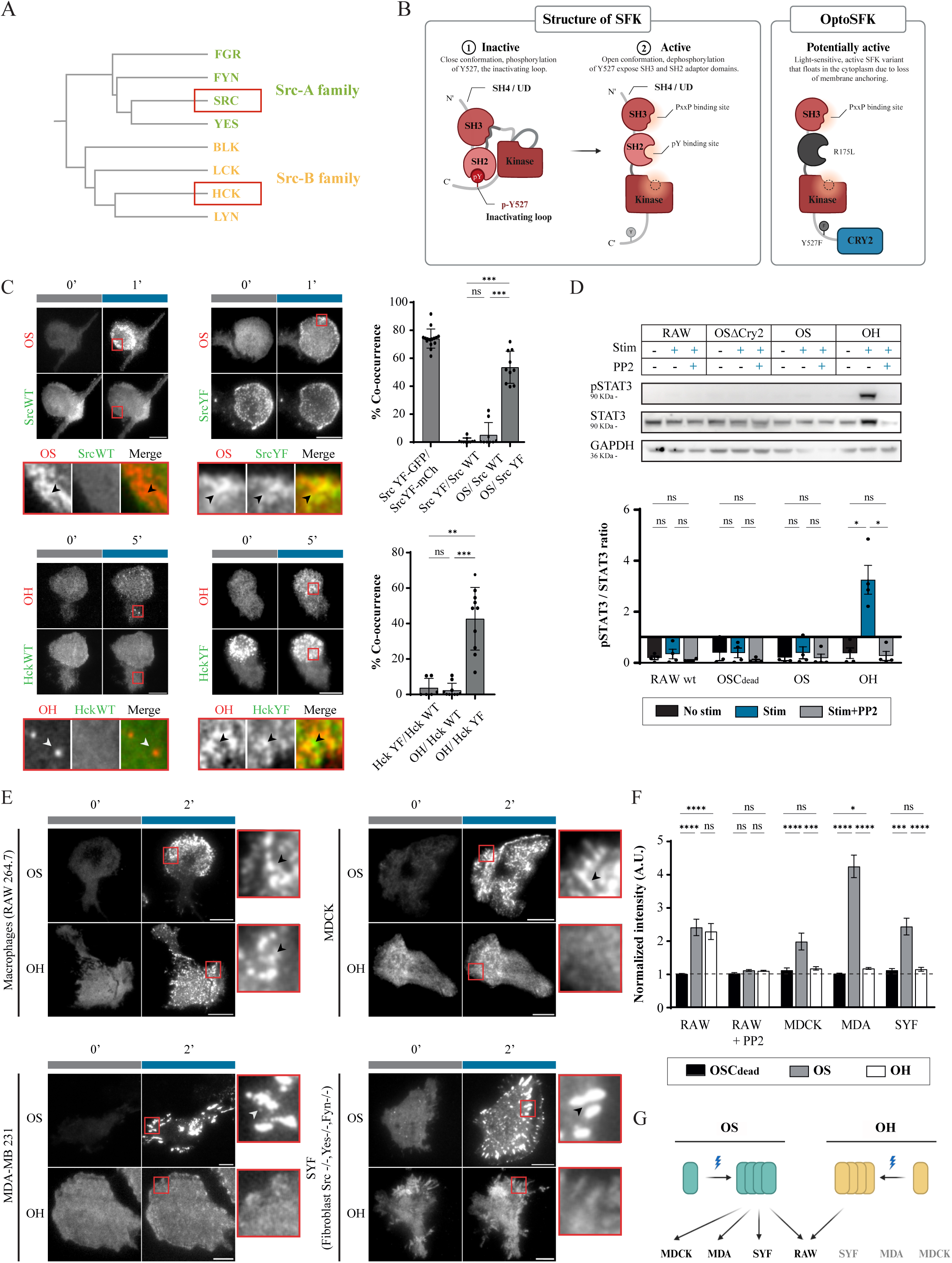
Optogenetic Src and Hck functionally mimic wild-type SFKs and recognize endogenous structures. **A.** Phylogenetic tree of Src Family Kinases (SFKs), depicting the Src-A and Src-B subfamilies based on protein sequence alignment. **B.** Schematic representation of the structure of the close conformation of endogenous SFKs and the engineered OptoSFKs (potentially active in the cytosol in the dark). **C.** Representative TIRF images of light-dependent recruitment of OSrc-mCherry (OS) to the plasma membrane of RAW 264.7 macrophages co-localizing with the constitutively active mutant Src Y527F-GFP (SrcYF) after 1 min of photo-stimulation but not with Src WT-GFP, while OHck-mCherry co-localized also with the constitutively active Hck Y522F-GFP (HckYF) after 5 minutes of photo-stimulation but not with Hck WT-GFP. The quantification of the percentage of co-occurrence confirmed that OS preferentially associates with active SrcYF structures; as well as OH with active HckYF structures. (N=3; > 10 cells/condition, mean ± SD. Statistical analysis with unpaired, Kruskal-Wallis test followed by Dunn’s post hoc test for multiple comparisons). **D.** Only light-dependent oligomerization of cytosolic OH specifically lead to STAT3 phosphorylation (measured by P-Stat3/total Stat3 ratio) upon 15 min of photo-stimulation in RAW 264.7 through a PP2-sensitive process (N= 4; normalized intensity ± SEM. Statistical test: unpaired, Kruskal-Wallis test followed by Dunn’s post hoc test for multiple comparisons). **E.** Representative TIRF images of the relocalization of OS or OH in response to blue light at the vicinity of the plasma membrane of different cell types (RAW 264.7, MDCK, MDA-MB 231 and SYF) expressing control constructs (OSCry2dead), after 2 min of photo-stimulation. **F.** Quantification of the normalized intensity measurements at 2 minutes post photo-stimulation revealed that OS relocalized in active sites across all cell types tested, whereas OH relocalization is restricted to RAW macrophages (N=10; 30 cells/condition; mean ± SEM; statistical test: unpaired, Kruskal-Wallis test followed by Dunn’s post hoc test for multiple comparisons). **G.** Schematic representation of OS and OH relocalization in different cellular lineage. Grey squares represent dark conditions (0 min), and blue squares indicate photo-stimulation. All TIRF experiments of photo-stimulation were performed at high stimulation frequency, 1 pulse every 10s (100 mHz). Statistical significance ns = no significant, *p<0.05, **p< 0.01, ***p< 0.001, ****p< 0.0001. Scale bar: 5 µm.

The redundancy between SRC and HCK can be largely attributed to their highly conserved structure and shared activation mechanism. Indeed, all SFK members share a canonical structure: an N-terminal SH4 domain responsible for membrane anchoring, followed by an intrinsically disordered unique domain (UD-the most divergent region among SFK members), an SH3 domain, an SH2 domain, and a C-terminal kinase domain. In the inactive state, SFKs are maintained in a closed conformation stabilized by two intramolecular interactions: one between the SH2 domain and a phosphorylated tyrosine in the C-terminal tail (Y527 in SRC), and a second between the SH3 domain and the internal proline-rich region (PRR, Fig 1B)^15,16^. Activation occurs when these interactions are disrupted, allowing the bi-lobal kinase domain to adopt an open conformation. This transition initiates a cascade of events marked by autophosphorylation of the activation loop and exposure of adaptor domains (SH3, PRR, SH2, and C-terminal tail) that engage in intermolecular interactions. Notably, activation promotes the formation of immobilized membrane-associated nanoclusters, dynamic assemblies of ∼6-15 SFK molecules confined within ∼80 nm regions, that serve as signaling hubs^17,18^. These nanoclusters, by locally concentrating SFKs, enhance signaling efficiency but it is not clear how these nm-organizations affect substrate selectivity.

It was proposed that the SH2/SH3-dependent adaptor functions of SFKs are key elements in substrate selectivity^19–21^. Besides activation, the SH2 domain is also known to guide SFKs localization to phosphotyrosine-rich complexes, influencing their subcellular localization^22^. The function of the SH3 domain in substrate selectivity is still not fully understood, despite its close coordination with the UD, which has been shown to play a role in substrates specificity^23^. Thereby, the combined contributions of these adaptor domains should allow the selection of specific and complex molecular environments where the kinase domain will be anchored transiently, phosphorylating accessible substrates to generate specific functional outputs. Till now, biochemical assays and phosphoproteomics have largely focused on the preferences of each kinase domain, although the influence of adaptor-mediated interactions on local signaling hubs is still unclear^20,24,25^.

Contrary to classical genetic loss-of-function approaches, optogenetics has emerged as a powerful strategy to manipulate intracellular signaling with high spatial and temporal precision^26^. To study SRC signaling in a dynamic context, we previously developed OptoSRC (OS) a photoactivable SRC kinase engineered using the CRY2 oligomerization module, enabling reversible nanoclustering of SRC with high kinetics (sec-min) and spatial (μm) control^27,28^. Unlike other optogenetic strategies that trigger full kinase activation by light-induced unfolding of the catalytic domain^29^, or those based on photoinhibition^30^, our approach selectively modulates SRC’s adaptor functions. Upon light stimulation, OS undergoes SH3-dependent binding to proline-rich proteins enriched at adhesive sites. By controlling the lateral mobility and diffusion of OS nanoclusters at the plasma membrane, we uncovered a key role for SRC’s spatial dynamics in shaping its pleiotrope signaling activity and inducing distinct cellular responses such as podosomes or lamellipodia formation^27^. Importantly, these light-driven probes operate within the same subcellular structures and kinetic range as endogenous SFK activity and show no leakiness in the dark, ensuring that the responses observed reflect physiologically relevant SRC-dependent signaling.

To revisit the concept of redundancy in cell signaling, optoHCK (OH) was engineered on OS’s model to control OS and OH adaptor functions via light-induced nanoclustering. Most surprisingly, OS and OH exhibited non-redundant and even opposite functions in macrophages. Light activation of OS promoted podosome formation and invasion, while that of OH disrupted them. These opposite effects were explained by their distinct SH3-dependent subcellular relocalization: OS accumulated at adhesive structures, whereas OH clustered to endocytic clathrin-positive structures, revealing a previously unrecognized role for HCK in regulating clathrin hotspots specialized in polarized clathrin-mediated endocytosis (CME) controlling directional macrophage migration. Modular optogenetic constructs were used to reintroduce redundant functions of OS within OH, showing that redundancy between SRC and HCK is supported by both their SH4 and SH2 domains. Therefore, rather than exhibiting strict redundancy, SRC and HCK appear to engage in a functional entanglement with shared activation dynamics but divergent spatial guidance, supporting a cooperative interplay wherein distinct and physically separated subcellular compartments (adhesive vs. endocytic). This work reframes kinase redundancy into a spatially encoded, modular signaling logic for cells to coordinate complex behaviors like adhesion, migration, and invasion.

## Results

### Activated OptoSRC (OS) and OptoHCK (OH) mimic key features of activated SRC and HCK

Dissecting redundancy between SRC and HCK requires determining whether these kinases can generate common cellular responses. However, this effort has been limited by the lack of tools enabling their real-time detection and discrimination, especially for HCK, which lacks specific biosensors or phosphor-specific antibodies (e.g. phospho-Tyr416-HCK). To overcome this, an optogenetic version of HCK, optoHCK (OH), has been built on the same model as the established optogenetic version of SRC, optoSRC (OS)^27^. The photosensitive CRY2-mCherry module was fused to a membrane anchor-deleted HCK mutant (ΔSH4 1-34) bearing a constitutively open conformation (Y522F) and a mutated SH2 domain preventing its binding to phosphor-tyrosine motifs (R171L, Fig. 1B). This minimal design retains the functional UD, SH3 domain, and kinase core, making the construct potentially active and freely cytoplasmic in dark conditions. We previously showed that OS has no off-target activity under dark conditions and mimics the physiological features of wild-type SRC function when activated by light^27^. Upon light-activation, it undergoes nanoclustering through CRY2 oligomerization^31^ and membrane recruitment enabling the precise spatial and temporal control of its kinase activity. To test the ability of OH to recapitulate endogenous HCK activity, we directly compared the subcellular localization of both optogenetic constructs (OS and OH) to their wild-type (WT) and constitutively active mutants fused to GFP (SRC-Y527F and HCK-Y522F, YF) at the vicinity of the plasma membrane by TIRF imaging. In the RAW 264.7 macrophage cell line, OS and OH exhibited diffuse cytoplasmic distribution in the dark, just like their respective WT counterparts (Fig 1C). Upon blue-light stimulation, both OS and OH nanoclusters transitioned from a diffuse cytoplasmic state to subcellular spots at the plasma membrane vicinity that precisely overlapped with the localization of their active YF counterparts (Fig 1C), with no evidence of off-pattern accumulation as quantified by a co-occurrence method. These results confirm that light-activated OS and OH faithfully recapitulate the spatial distribution of active endogenous SRC and HCK.

Beyond relocalization, light activation of OS and OH triggered their kinase activity, as shown by an increase in protein phosphorylation that was strongly suppressed by the SFK inhibitor PP2 (data not shown). Notably, OH induced a stronger global phosphorylation signal than OS, and uniquely led to STAT3 phosphorylation within just 15 minutes of light stimulation in RAW cells (Fig. 1D), which aligns with the established role of HCK in regulating STAT3 signaling in myeloid cells^32–34^. Therefore, these data support the physiological relevance of the optoHCK design and point toward a specific functional signature for OH.

Since HCK is mostly expressed in myeloid lineage, the physiological aspects of OH relocalization in response to its light-dependent nanoclustering were challenged by comparing it in myeloid and non-myeloid cell types (fibroblasts, epithelial and cancer cells, movie S1 and S2). Importantly, OH relocalization occurred only in cells that endogenously express HCK (RAW 264.7 Fig.1E, and primary bone marrow-derived macrophages BMDM, Fig S1D), and was absent in non-myeloid cells (MDCK, MDA-MB 231, SYF fibroblasts Src-/-Fyn-/-Yes-/-; Fig.1E-F), with only minimal relocalization observed in HeLa and MEFs (data not shown). In contrast, activated OS consistently localized to adhesion structures (podosomes or focal adhesion) across a broad range of cell types, consistent with the ubiquitous expression of SRC. In RAW cells, activated OH accumulated in subcellular compartments that appeared different from those targeted by OS which were more reminiscent of podosomes clusters (Fig.1E). This shows that the relocalization of OH nanoclusters can only occur in specific cellular environments, most likely where HCK’s SH3 domain encounters compatible proline-rich binding partners.

Together, these results demonstrate that light-induced nanoclustering of OS and OH mimic the physiological activation of these kinases and arise in different subcellular compartments. While activated OS targets adhesion-related compartments correlated with the ubiquitous expression of SRC, activated OH relocalization and signaling are restricted to myeloid cells, where its SH3 domain is most likely encountering compatible partners (Fig 1G).

### Activated OH is not functionally redundant with OS since it induces opposite adhesives responses in macrophages

To challenge the potential functional redundancy between OS and OH, we examined whether HCK activation could mimic the well-characterized effects of SRC activation on podosome dynamics (Destaing et al., 2008). Using TIRF microscopy, RAW 264.7 macrophages stably co-expressing either OS or OH together with LifeAct-GFP were subjected to high-frequency cyclic blue-light stimulation (100 mHz) to assess podosome organization.

Activated OS rapidly (within 2 min) accumulated at podosomes, as visualized by strong co-occurrence with Lifeact-GFP. Consistent with SRC’s established role in orchestrating podosome dynamics, high-frequency blue-light stimulation of OS (33-100 mHz, during 10 min) triggered robust podosome assembly, including the formation of polarized podosome rings at one cell edge (Fig. 2A, movie S3). This was induced by its CRY2-dependent nanoclustering as OS mutated on CRY2 (OS-Cry2Dead, not able to bind FAD and no nanoclustering under blue light) did not stimulate podosome formation (Fig.2A). In contrast, OH activation showed a clearly distinct subcellular localization pattern, with minimal overlap with podosome-rich zones (Fig. 2A-B, movie S4). Moreover, under the same high frequency stimulation conditions as for OS, this activation of OH away from podosomes led to a rapid and graded disassembly of podosomes (Fig. 2A-B). This result was unexpected as SRC and HCK are recognized as the most redundant SFKs in the myeloid lineage^35^.

**Figure 2:**
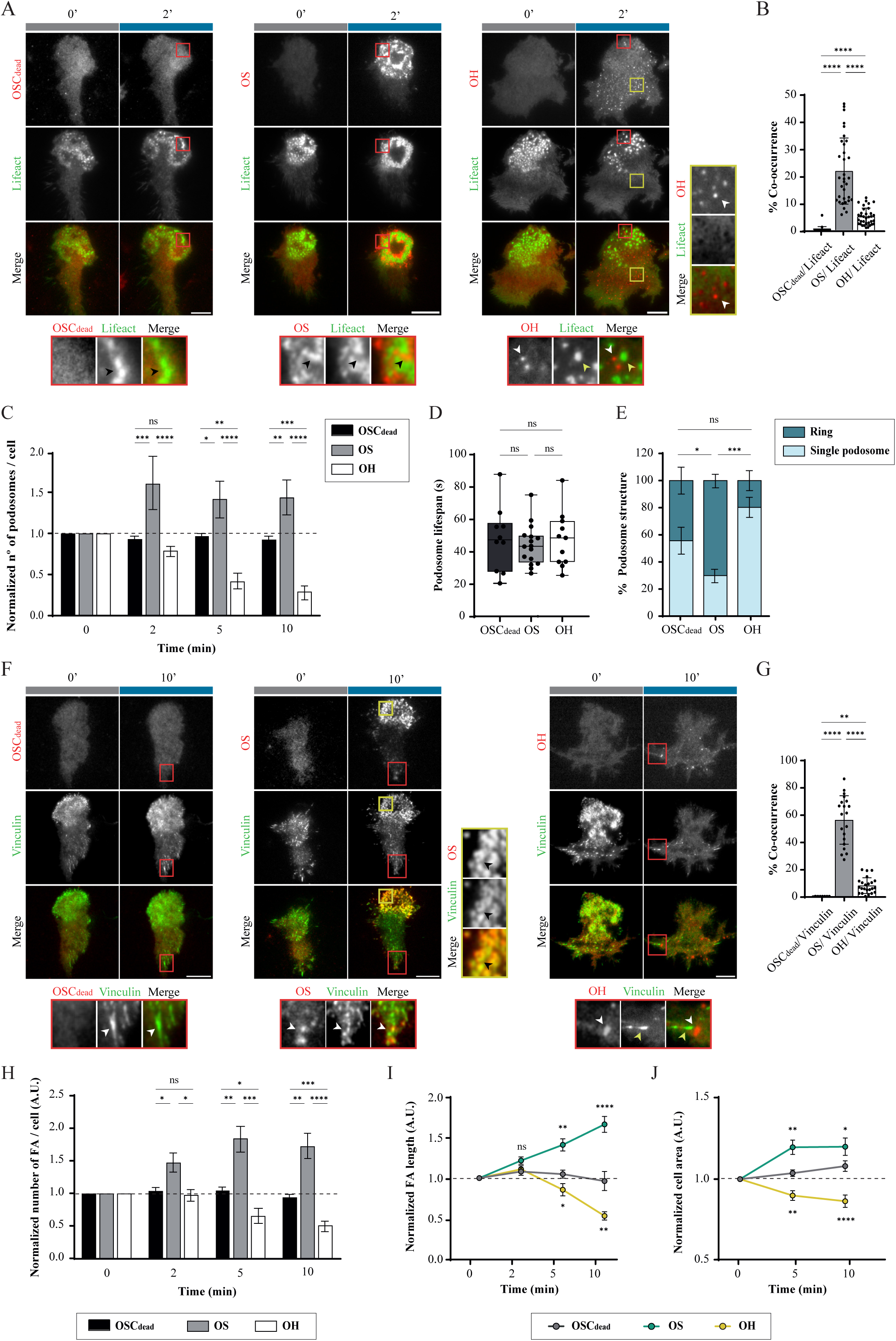
Light-activation OS induces podosome and focal adhesion formation while OH disrupts them both. **A.** Representative TIRF images show light-activation of OS lead to its specific relocalization in podosomes (followed by Lifeact-GFP) of RAW 264.7 macrophages and increases the formation of larger podosomes rings only in 2 min, while activated OH are not relocalizing to podosomes but rather induced their disorganization. **B.** Quantification of co-occurrence between OSCry2dead, OS, or OH and Lifeact-GFP confirmed that only OS specifically relocalizes to podosomes. (N=3; 20-30 cells/condition; mean ± SD; statistical test: unpaired, Kruskal-Wallis test with Dunn’s post hoc test. **C.** Quantification of normalized podosome proportion after 10 min of photo-stimulation showed that OS promotes podosome formation, while OH disrupts them. (N=4; 20 cells/condition; mean ± SEM; statistical test: unpaired, 2-way ANOVA with Tukey’s post hoc test). **D.** Podosome lifespan per cell remains unchanged during 10 min of photo-stimulation. (N=3; 10-15 cells/condition; mean ± SEM; statistical test: unpaired, 2-way ANOVA with Tukey’s post hoc test). **E.** Quantification of the different podosome organization showed that OS activation significantly promotes ring-like structures, whereas OH activation leads to dispersed single podosomes. (N=5; 30-35 cells/condition; mean ± SEM; statistical test: unpaired, 2-way ANOVA with Tukey’s post hoc test). **F.** Representative TIRF images show light-activation of OS lead to its specific relocalization in focal adhesion (followed by Vinculin-GFP, 10 min) of RAW 264.7 macrophages, while activated OH is not relocalized in these other adhesive structures. **G.** Quantification of co-occurrence between OSCry2dead, OS, or OH and Vinculin-GFP positive structures confirmed that only OS specifically relocalizes to focal adhesions. (N=5; 20-30 cells/condition; mean ± SD; statistical test: unpaired, Kruskal-Wallis test with Dunn’s post hoc test). **H.** Quantification of the normalized variation of the number of focal adhesion (FA) over 10 min of photoactivation showed that OS increases their formation, while OH disorganizes FAs (N=3; 14 cells/condition; mean ± SEM; statistical test: unpaired, 2-way ANOVA with Tukey’s post hoc test). **I.** Normalized FA length shows that OS increases significantly FA size, whereas OH reduces it over time of photoactivation (O-2-5-10 min) (N=3; > 50 focal adhesion/condition; mean ± SEM; statistical test: unpaired, 2-way ANOVA with Tukey’s post hoc test). **J.** Normalized cell area over 10 min of photo-stimulation shows that OS induces cell spreading, while OH causes cell retraction (N=3; > 30 cells/condition; mean ± SEM; statistical test: unpaired, 2-way ANOVA with Tukey’s post hoc test). Grey squares represent dark conditions (0 min), and blue squares indicate photo-stimulation. All TIRF experiments of photo-stimulation were performed at high stimulation frequency, 1 pulse every 10s (100 mHz). Statistical significance ns = no significant, *p<0.05, **p< 0.01, ***p< 0.001, ****p< 0.0001. Scale bar: 5 µm.

OS increased podosome number per cell by ∼50%, whereas OH disorganized them by more than 50% (Fig.2C). These changes in podosome organization were not associated with a change in their life-span (Fig.2D), indicating that OS and OH were mostly controlling podosome assembly. In addition, OS activation induced clustering of podosomes into ring-like metastructures, while OH disorganized them into individual podosomes (Fig.2E).

The actions of OS and OH were also analyzed on focal adhesions (vinculin-GFP positive compartments), a secondary adhesion structure also present in macrophages. Interestingly, contrary to activated OS, activated OH did not localize to vinculin-positive focal adhesions at the cell rear (Fig. 2F-G). As for podosomes, OS activation significantly increased (∼50%) focal adhesion number and size (Fig. 2H-I), along with enhanced cell spreading (Fig. 2J), effects that were again absent with OS-Cry2Dead (Fig. 2H-I). Conversely, OH activation reduced focal adhesion size and number, and decreased spreading (Fig. 2H-J). In non-myeloid cells, MDA-MB-231, OH activation had no impact on focal adhesions, consistent with its lack of OH relocalization. In MEFs, however, where OH did relocalize, it induced a marked reduction in focal adhesion number and spreading, reinforcing the causal effect between relocalization of activated OH at the membrane and potential effects on cell spreading and adhesion in different cell types.

Together, these findings reveal not only distinct spatial localizations but also strikingly opposite functions for these two kinases in macrophages despite their similarity and proposed redundancy. The divergent localizations and functions of OS and OH are most likely driven by their SH3-mediated interactions with specific partners that drive their distinct subcellular niches.

### Activated OptoHCK is specifically relocalized in a subset of clathrin structures

To elucidate the specific functions of OH in macrophages, the identity of the subcellular structures where active OH relocalizes in response to light was investigated. As previously noted, HCK has been difficult to study due to the lack of specific tools with no cross-reactivity with other SFKs. As a result, most insights rely on overexpression studies, placing HCK in various compartments, including podosomes, unconventional lysosomes, and the plasma membrane^11,36,37^. However, overexpression of HCK -and other SKFs-can drastically perturb cellular homeostasis, including inflammatory responses, impair phagocytosis, and disrupt autophagy. On the contrary, the light-inducible version of HCK (OH) allows for a transient temporal control of its activity with no background signal in the dark, therefore minimal side-effects.

To further investigate the nature of the spots where OH relocalized, their spatial distribution and dynamics were studied in polarized macrophages. In our culture conditions, RAW macrophages exhibit a front-rear polarity: a front region defined by a lamellipodium enriched in podosome clusters or rings, and an elongated rear region where podosomes are scarce (Fig. 3A). Live TIRF microscopy revealed a striking asymmetry: long-lived, stable OH spots accumulated predominantly at the rear (Fig. 3A, zoom and kymogram rear-black arrows), whereas transient, short-lived spots were preferentially formed at the front (Fig. 3A, zoom and kymogram front-black arrows). Kymograph analysis (Fig.3A) and quantification over time confirmed this polarity, with a significantly higher persistence and density for OH spots at the rear region of macrophages (Fig. 3B). To analyze the underlying motion of these structures at high throughput level, a particle tracking analysis was applied to thousands of individual OH spots. Two distinct dynamic populations appeared: highly mobile spots mostly at the front, characterized by extended track lengths over brief lifespans (less than 5 min after light stimulation), and long-lasting and immobile spots at the rear (more than 10 min after light stimulation, Fig. 3C). Notably, rear-enriched OH spots included a population with ∼50% higher fluorescence intensity, suggesting increased local accumulation of activated OH molecules per spot. The ability of OH spots to be either immobilized or move in a plane at the vicinity of the basal plasma membrane suggests that OH spots are not passive aggregates but resemble vesicle-coated structures involved in membrane trafficking segregated in polarized macrophages.

**Figure 3:**
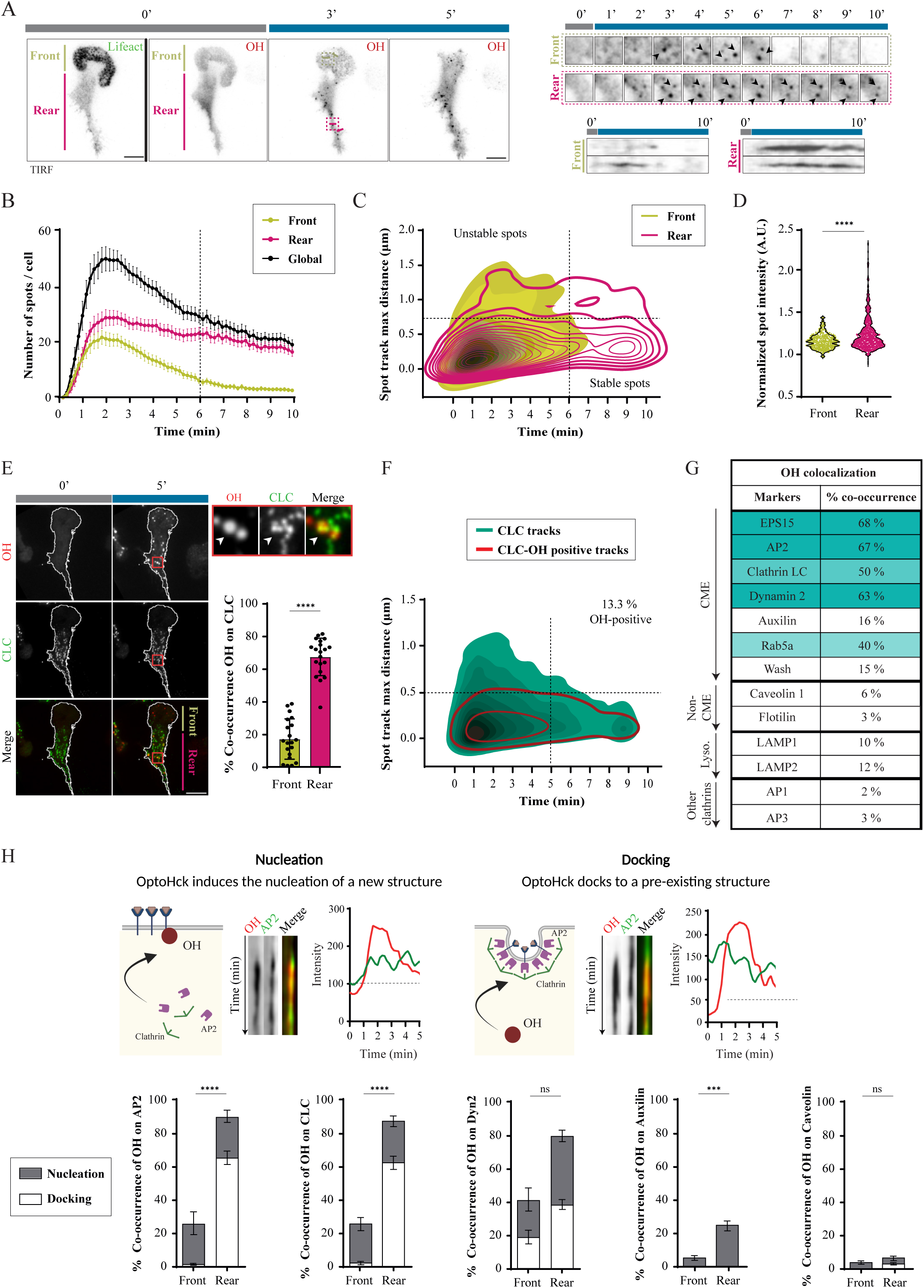
OH preferentially recruits to the rear of polarized macrophages, associating with long-lived Clathrin-Mediated Endocytosis (CME) sites. **A.** Representative TIRF images showed that the relocalization of activated OH in polarized RAW 264.7 macrophages spatially defined in two regions: the front region which is characterized by the presence of podosomes (Lifeact-GFP, pistachio) and the rear region corresponding to the rest of the cell body (fuchsia). Representative zoomed TIRF images and kymographs show the high stability of OH spots (black arrows) in the rear of the cell in comparison to transient behaviors in the front. **B.** Quantification of OH spots per cell shows a higher accumulation of OH at the rear of the cell than in the front. (N=5; 60 cells/condition; mean ± SEM). **C.** Quantification of both maximal distance and spot lifespan from tracking analysis of OH spots reveals two kinetically distinct behaviors (unstable and stable spots) in RAW 264.7 macrophages over 10 min of photo-stimulation. OH spots appeared more stable when observed in the rear region of the cell (fuchsia lines) while in the front (pistachio; N=4; 30 cells/condition; data are presented as KDE plots). **D.** Normalized OH spot intensity analysis shows that OH spots at the rear are significantly brighter in the rear than in the front region. (N=5; 10-15 cells/condition; statistical test: unpaired, Mann-Whiteny test). **E.** Representative TIRF images of activated OH relocalization in RAW 264.7 macrophages show its co-localization with clathrin structures (CLC-GFP, 5 min post-stimulation). Quantification of co-occurrence between activated spots and CLC-GFP reveals spatial differences, with significantly higher colocalization at the rear region. (N=3; 20 cells/condition; mean ± SD. Statistical test: unpaired, *t*-test). **F.** Tracking of clathrin structures (CLC-GFP) over 10 min of OH activation reveals two different clathrin populations (green): unstable structures characterized by short lifespan able to travel on long distance, and stable structures immobile over many minutes. OH selectively associates with stable clathrin structures (red) and only in 13,3% of the case. (N=5; 15 cells/condition; data are presented as KDE plots). **G.** Quantification of the co-occurrence of activated OH spots with numerous endocytic and vesicle trafficking markers confirms its specific association with Clathrin-Mediated Endocytosis (CME) process. (N=3; 5-20 cells/condition). **H.** Schematic representation of OH role in CME that can either nucleate new CME sites when being relocalized at the vicinity of the plasma membrane in response to light, or dock onto pre-existing ones. Both behaviors have been analyzed from kymograph analysis followed by RGB profile analysis. For nucleation, OH spots are associated with recruitment of any endocytic marker (AP2-, CLC-, Dyn2- or auxillin-GFP) while OH spots accumulate on already existing endocytic structures in the case of docking. Manual quantification of OH co-occurrence with AP2-, CLC-, Dyn2-, Auxilin- and Caveolin-GFP were performed both at the front and rear of polarized macrophages. Grey bars indicate OH nucleation (marker recruitment), while white bars indicate OH docking to pre-existing structures. (N=3; 10-20 cells/condition; mean ± SEM; statistical test: unpaired, 2-way ANOVA test). Grey squares represent dark conditions (0 min), and blue squares indicate photo-stimulation (A-G). All TIRF experiments of photo-stimulation were performed at high stimulation frequency, 1 pulse every 10s (100 mHz). Statistical significance: ns = no significant, *p<0.05, **p< 0.01, ***p< 0.001, ****p< 0.0001. Scale bar: 5 µm.

To determine the identity of these structures, we investigated their association with endocytic structures. In polarized RAW macrophages co-expressing OH and Clathrin Light Chain-GFP (CLC-GFP), light-activated OH co-localized with clathrin predominantly at the rear (Fig. 3E, movie S5), but only with a specific subset (13.3%) of the total clathrin-coated structures (CCS, Fig. 3F). These CCS were immobile and could be long-lived at the plasma membrane, pointing to a specialized subpopulation of CCS. OS, in contrast, showed minimal clathrin association (movie S6). Kernel density estimation (KDE) plots revealed that only OH increased the population of stable CCS, highlighting the active effect of its targeting to this subpopulation of CCS.

Activated OH was essentially associated with canonical clathrin-mediated endocytosis (CME) markers, including Eps15-GFP, AP2-GFP, dynamin2-GFP, and even present on early endosomal Rab5a-GFP vesicles that follow clathrin-mediated internalization (Fig. 3G). Despite the imaging challenge posed by rapid turnover of some markers, OH was also detected to be transiently associated with WASH-GFP (actin regulator) and auxilin-GFP (clathrin uncoating factor).

Conversely, OH did not colocalize with caveolin-GFP (caveolae-mediated endocytosis), flotillin-GFP (raft-mediated endocytosis), or LAMP1/2-GFP (lysosomes). Activated OH also did not associate with AP1, which marks the trans-Golgi network and early endosomes, nor with AP3, which is involved in endolysosomal trafficking, reinforcing its specificity for plasma membrane AP2-positive clathrin structures (Fig.3H). This selective affinity for a specific subset of CCS (CLC- and AP2- positive long-lasting CCS in macrophage) likely reflects OH’s selective engagement with a subset of specific proline-rich partners in a subpopulation of CCS enriched in macrophages, absent or less abundant in non-myeloid cells such as MDCK or MDA-MB 231. To determine whether OH nanoclusters nucleate new CCS or dock onto pre-existing ones, their dynamics of localization followed by TIRF imaging were analyzed through two-color kymograph analysis (Fig. 3H). Most frequently, activated OH recruites to pre-existing, long-lasting CCS at the rear of macrophages, indicating a docking mechanism (Fig. 3I). In approximately 20% of events, OH nanoclusters preceded clathrin recruitment, suggesting it can also participate in nucleation events (Fig. 3I). Similar dynamics were observed with AP2-GFP. Co-occurrence analysis with dynamin2-GFP, which is enriched during both maturation and fission of CCS, showed relocalization of activated OH before and after the dynamin2 recruitment (50% for both behaviors) at the front and rear regions of polarized macrophages. Supporting this, markers of late-stage CME (auxilin and Rab5a) appeared mostly after activated OH (Fig. 3I).

Thus, light-induced OH nanoclustering reveals a highly specific interaction with a distinct, stable subpopulation of CCS, predominantly found in macrophages. These findings highlight the spatial and functional heterogeneity of CCS in macrophages and position OH as a unique tool for dissecting specialized trafficking pathways associated with HCK function.

### Activation of optoHCK reveals and modulates clathrin endocytosis hotspots in macrophages

This specific interaction between activated OH and clathrin-coated structures (CCS) prompted us to investigate its impact on clathrin organization and the dynamics of this CCS subtype in macrophages. Notably, light activation of OH -but not OS- induced a rapid and significant increase in the number of CLC-GFP-positive structures at the basal plasma membrane, within just a few minutes (Fig. 4A). This increase could be due to the formation of new CCS, an inhibition of fission in existing CCS, an increase of fission leading to moving CCS or a combination of them. High frequency TIRF imaging revealed that OH nanoclusters did not accumulate on single clathrin-coated pits (CCPs) but rather localized to areas surrounded by multiple and dynamic clathrin puncta (red arrows Fig. 4B). These areas appeared to act as nucleation or docking platforms where CCS were either newly formed or spatially organized. To assess how activated OH influences CCS dynamics, intensity analysis of two-colors kymograph of OH-mCherry and CLC-GFP was performed on polarized macrophages. At the cell front, short-lived OH accumulations were associated with the nucleation of transient CCS (first panel-Front Fig. 4C). At the cell rear, rhythmic fluctuations in CLC-GFP intensity were observed suggesting repeated cycles of CCP formation and fission associated with long-lasting OH spots (second panel-Rear Fig. 4C). These events were specific to OH, since traces negative for OH did not reveal the same rhythmic fluctuations. Quantitative analysis of CLC-GFP intensity fluctuation showed that OH-enriched regions at the rear of macrophages displayed significantly higher pulsatility (more peaks per unit of time), compared to OH-negative areas, even within the same cell. Moreover, these fluctuations were more homogenous, as indicated by a lower variation between pulse amplitudes across time (Fig. 4D). This suggests that OH nanoclusters are localized to highly dynamic and potentially coordinate endocytic hotspots by enhancing or stabilizing cycles of CCS maturation and fission. Importantly, the occurrence of these rhythmic, pulsatile CCS was not only dependent on OH activation itself. In CCS negative for OH, rhythmic CCS dynamics could still be detected in macrophages (Fig.4D), confirming that OH preferentially targets and modulates a pre-existing subpopulation of active, long-lived CCS hotspots rather than inducing them de novo or artifactually.

**Figure 4:**
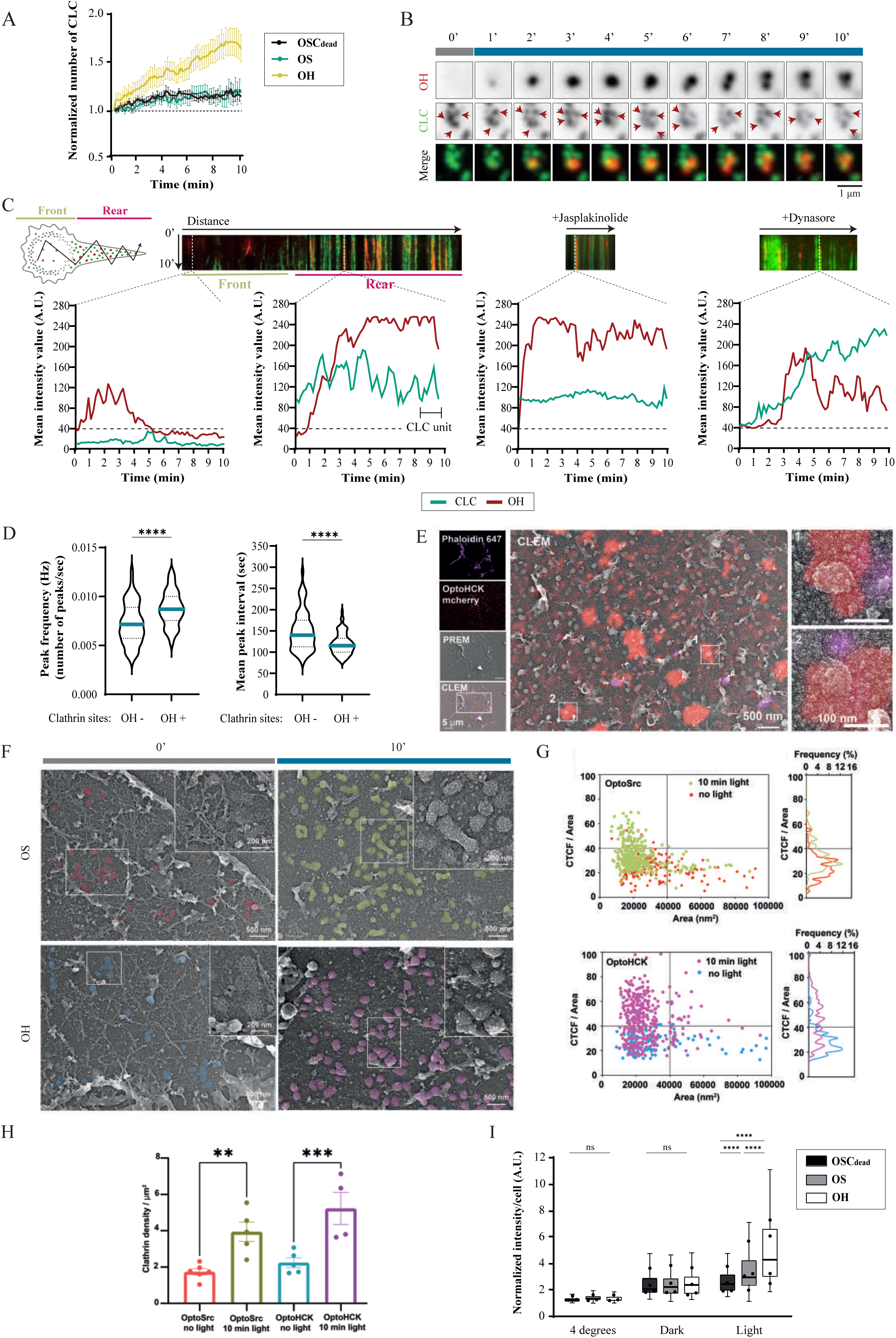
Activated OH revealed the existence of stable clathrin hotspots and allowed to increase CME in macrophages. **A.** Quantification of the normalized number of CLC-GFP spots per cell over 10 minutes of photo-stimulation shows that only OH increases clathrin accumulation over time. (N=3; *n* > 15 cells/condition; mean ± SEM). **B.** Cropped representative time-lapse TIRF images show the behaviors of CLC-GFP during the formation of an OH spot at the rear region. Multiple clathrin structures (CLC-GFP, red arrows) dynamically emerged at the same site over time, suggesting that OH defines stable clathrin hotspots. **C.** Representative kymographs of polarized RAW 264.7 macrophages co-expressing OH and CLC-GFP and photoactivated reveal spatial differences in the dynamics of clathrin structures. RGB profiles of the kymograph on OH spots at the rear region show multiple bursts of CLC-GFP over time, defining a, stable clathrin hotspot behavior. At the front, transient OH colocalizes with transient single clathrin burst. Jasplakinolide pre-treatment disrupts hotspot behavior in OH spots by preventing the repeated recruitment of clathrin to the same location. Inhibition of the fission enzyme Dynamin by dynasore treatment generates accumulation of clathrin on OH sites. **D.** Quantification of peak frequency and mean peak variation of CLC-GFP intensity associated (OH +) or not (OH −) with OH spots present in the rear region of polarized macrophages. OH increases the homogeneity of CLC-GFP pulsing and thus hotspots behaviors. (N=3; 9 cells per condition with around CLC-GFP 1000 traces analyzed; data are represented as violin plots; statistical analysis: unpaired, Mann-Whitney test). **E.** Correlative fluorescence microscopy (CLEM) and PREM imaging of RAW 264.7 macrophages expressing OH-mCherry after 10 minutes of photoactivation shows nanoscale localization of OH (red) surrounding clathrin-coated pits seen by PREM. Phalloidin (magenta) was used to visualize actin structures. Insets highlight OH accumulation near CCSs. Scale bars: 5 µm, 2 µm, 0.5 µm, and 0.1 µm (insets). **F.** Representative high magnification platinum replica electron microscopy (PREM) images of unroofed RAW 264.7 macrophages show clathrin structures under different conditions. Macrophages expressing OS or OH were imaged without photoactivation or after 10 minutes of light stimulation. Clathrin structures are pseudo-colored to highlight each condition. Insets, displayed at equal scale, show magnified views of representative CCSs. Scale bars: 5 µm, 0.5 µm, and 0.2 µm (insets). **G.** Morphometric analysis of individual CCSs in OS- and OH-expressing macrophages in the dark or after 10 minutes of photoactivation, displayed as dot plots of curvatures (gray-level intensity) versus area. Each structure was manually segmented based on its honeycomb morphology, and the percentage distribution across curvature/size quadrants is indicated for each condition. This reveals that OH activation induces higher curvature of the clathrin-coated pits. **H.** Quantification of CCS density (mean density per µm^2^) reveals a significant increase after photoactivation of OH compared to the dark condition, while OS photoactivation induces a moderate increase. (N=3; 150-320 clathrin structures analyzed; mean ± SEM; statistical analysis unpaired, non-parametric one-way ANOVA). **I.** Quantification of normalized transferrin intensity per cell-endocytosis assay-in RAW 264.7 macrophages expressing OSCry2dead, OS, or OH under different conditions (4 °C, no light, and 10 min photo-stimulation) show that OH induced a strong increases of CME just after few min of photoactivation (N=4; > 30 cells/condition; Box-and-whisker plots (Tukey style) display the distribution, with overlaid points representing the mean of each replicate; statistical analysis: unpaired, non-parametric one-way ANOVA). Grey squares represent dark conditions (0 min), and blue squares indicate photo-stimulation. All TIRF experiments of photo-stimulation were performed at high stimulation frequency, 1 pulse every 10s (100 mHz). Statistical significance: ns = no significant, *p<0.05, **p< 0.01, ***p< 0.001, ****p< 0.0001. Scale bar: 2 µm (B).

Treatment with jasplakinolide, an inhibitor of actin polymerization that blocks CCS maturation membrane fission, completely abolished this pulsatile behavior (Fig. 4C third panel). This confirms that the observed intensity fluctuations represent bona fide complete endocytic events (from nucleation to efficient fission). Importantly, OH nanoclusters remained colocalized with CLC-GFP even in the presence of jasplakinolide, indicating that OH relocalization occurs upstream of or independently from vesicle scission.

In contrast, treatment with dynasore, which inhibits dynamin-2 that is implicated mostly in CCS fission but also in its maturation^38,39^, did not completely block CLC-GFP pulsatile activity but led to its limited accumulation at the membrane (Fig.4C fourth). In addition, dynasore limited OH recruitment, without affecting its colocalization with CLC-GFP, suggesting that dynamin2 activity is required for efficient OH enrichment at clathrin hotspots, potentially by enabling a mature structural or signaling context favorable for OH docking. Altogether, these elements support that activated OH relocalized to clathrin hotspots and regulates their pulsatile activity, suggesting that they are intense hotspots of functional CME.

The ultrastructure of clathrin-coated structures was analyzed using correlative light and electron microscopy (CLEM). Combining super-resolution to platinum replica EM (PREM) reinforced the link between activated OH nanoclusters and CCPs sites (Fig.4E). PREM of unroofed adherent macrophages was performed in order to determine if these CCS hotspots were clathrin lattice, as sometimes described in other systems^40^. However, instead they consisted of densely packed individual CCPs (Fig. 4F) which density was significantly increased upon OH activation (Fig. 4H). Strikingly, OH also enhanced the membrane curvature of these CCPs (Fig. 4G) as quantified by the ratio of the fitting pit contours on measured curvature parameters (CTCF). Although OS activation induced a mild increase in CCP density, it did not alter pit curvature, likely reflecting potential indirect effects perhaps via partial cross-activation of endogenous HCK by OS.

Functionally, the OH-dependent increased CCS density and rhythmic fluctuation of CCP into these hotspots should affect CME efficiency. Transferrin uptake assays confirmed that OH activation nearly doubled its internalization rates (Fig. 4I), underscoring OH ability to both recognize and boost the function of clathrin hotspots in macrophages.

Altogether, these results not only reveal a distinct, previously uncharacterized subpopulation of CCS composed of dynamic clathrin hotspots but they demonstrate that HCK actively enhances their endocytic function in macrophages. Beyond shedding light on a functional heterogeneity among SFK, these results demonstrate that the optogenetic control of SFK nanoclustering can serve as a powerful tool to uncover new cell-type-specific features and regulatory modules in subcellular organization.

### Functional synergy of SRC and HCK in macrophages functions

As shown above, at high-frequency light stimulation (33-100 mHz, Fig.2), optogenetic activation of SRC and HCK revealed distinct localization patterns and antagonistic effects on podosome formation. However, since genetic studies have suggested that SRC and HCK have highly redundant cellular function^6^, we examined how OS and OH affect cell migration and invasion, two cellular outcomes known to be regulated by podosomes and SFKs. This approach aimed to dissect not only short-term signaling dynamics, but also how sustained activation of SFK shapes long-term cellular behaviors.

Slow cyclic light (4 mHz) activation was applied to mimic milder and sustained SFK activation over time, ensuring periodic reactivation of the kinase to generate a lower plateau of signaling activity than with 100mHz activation^41^. The impact of OS and OH activation on macrophages migration and invasion was monitored 3 hours before and during slow cyclic blue light stimulation (Fig.5A-B). OS activation led to a marked immobilization of macrophages, whereas OH activation significantly enhanced migration speed (Fig. 5B-C) and directionality, indicating a shift toward more persistent migration (Fig. 5D). Quantification of hundreds of cells per condition confirmed that OS-expressing cells showed reduced migration distance and speed, while OH-expressing cells increased their migration by nearly 50%. To determine whether this modulation extended to invasion, we used Boyden chambers coated with Matrigel to assess extracellular matrix (ECM) invasion capacity (coupling migration and ECM degradation). Remarkably, differently from migration, only OS activation enhanced macrophage invasion, while OH activation had no significant effect (Fig. 5E). Notably, this lower plateau of OH activation did not alter the number of podosomes per cell over time in this condition of activation (Fig.5F). Thus, while podosomes organization remained unchanged, SRC activity promoted pro-invasive behavior through podosome activation, whereas HCK promoted pro-migratory behavior, likely via mechanisms involving podosome turn-over and endocytic regulation. Together, these findings reveal that SRC and HCK activities in macrophages could not be merely antagonistic but could be functionally specialized. HCK promotes migration through spatially confined, regulated endocytic hotspots, while SRC supports matrix invasion, possibly through global podosomes activation and persistent adhesion. Importantly, these functional compartments -clathrin hotspots and podosomes- are spatially distinct within the macrophage, typically separated by several microns, yet they functionally cooperate to fine-tune cellular responses across different timescales.

**Figure 5:**
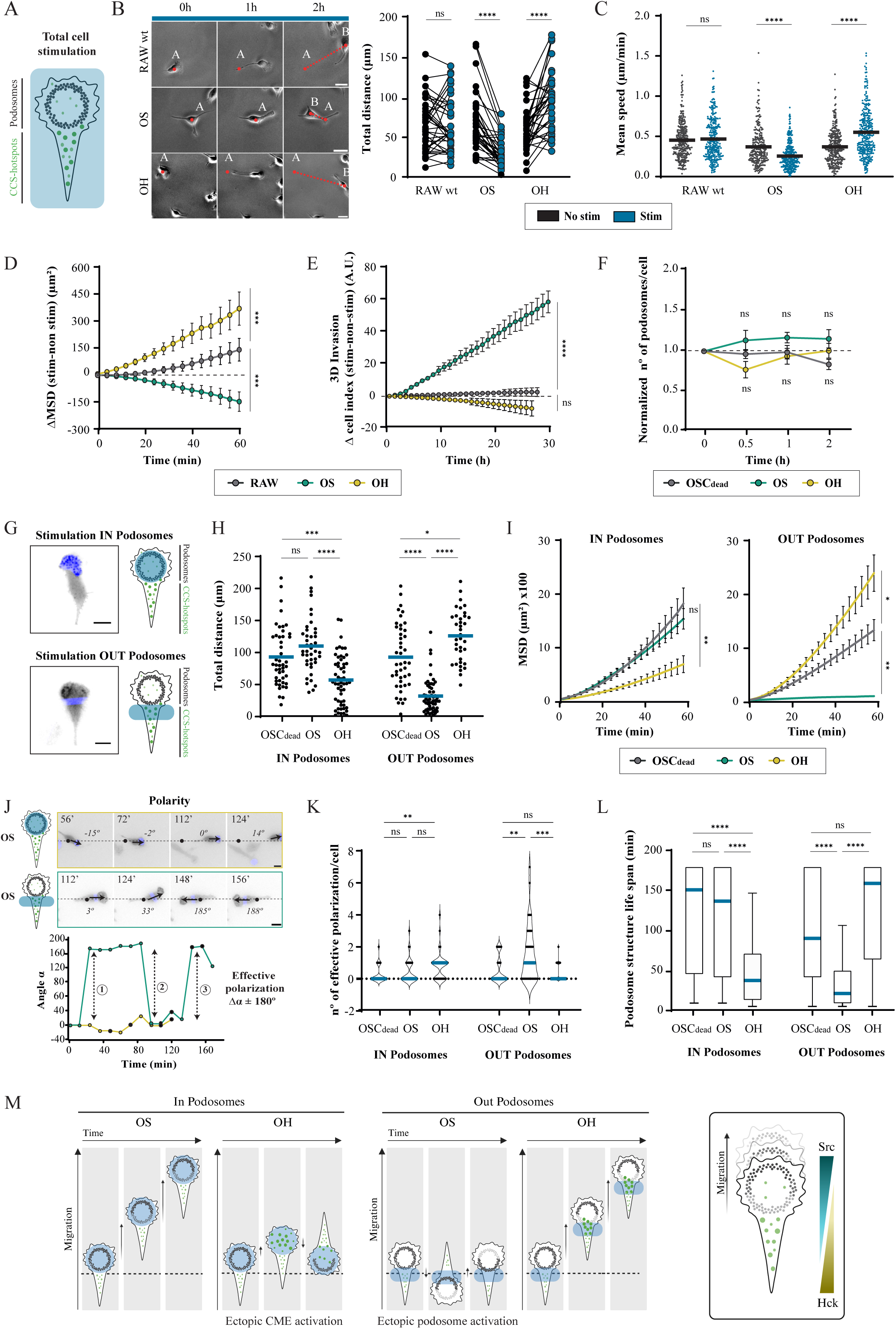
Spatially controlled activation of OS and OH synergize to regulate macrophage migration and polarity. **A.** Schematic representation of macrophages fully stimulated spatially with podosomes (grey) at the front and CCS (green) at the rear. **B.** Representative transmission images of fully photoactivated cells showing 2D migration after global illumination of RAW wild-type cells, expressing stably OS or OH. Quantification of the total distance traveled by RAW WT, OS or OH before and after photoactivation over 3 h. Most of the OS cells exhibit reduced migration, whereas OH macrophages show enhanced migration. (N=4; > 30 cells/condition; statistical test: unpaired, Mann-Whitney test). **C.** Quantification of the mean speed of populations of macrophage with or without photo-activation. Again, OS-expressing macrophages show a global reduction in motility, while OH-expressing macrophages display an overall increase in migration speed. (N=4; 300 cells/condition; lines represent the mean speed of the population; statistical test: unpaired, Mann-Whitney test). **D.** Difference in Mean Square Displacement (MSD) before and after photo-stimulation shows that OH-expressing macrophages exhibit significantly more effective and directional migration, while OS-expressing macrophages lose directionality in comparison to RAW WT cells. (N=3; 30 cells/position; mean ± SEM; statistical test: unpaired, one-way ANOVA with Tukey’s post hoc test). **E.** Quantification of 3D invasion (shown as the difference in cell index) through Matrigel, using xCELLigence, reveals that only OS enhances invasion capacity. (N=3; mean ± SEM; statistical test: unpaired, one-way ANOVA with Tukey’s post hoc test). **F.** Quantification of normalized podosome count per cell after 2 h of photo-activation at low-frequency (4mHz) shows that neither OS nor OH significantly change the number of podosomes per cell. (N=3; 10-20 cells/condition; mean ± SEM; statistical test: unpaired, 2-way ANOVA with Tukey’s post hoc test). **G.** Schematic representation of the strategy of spatially constrained photo-activation. Blue circle or square indicates the region of interest: IN where photo-stimulation was applied relative to podosome location, or OUT representing photo-stimulation outside podosome location. **H-I.** Quantification of 2D migration and MSD of RAW expressing OSCry2dead, OS, or OH under localized photo-stimulation (IN vs. OUT of podosomes). Activation of OS within podosomes enhances migration and maintains directionality, whereas stimulation outside podosomes impairs both. In contrast, OH stimulation within podosomes reduces migration and directionality, while stimulation outside podosomes promotes them (N=3; 40-50 cells/condition; statistical tests: unpaired, Kuskal-Wallis with Tukey’s post hoc test for H; unpaired, one-way ANOVA with Tukey’s post hoc test for I). **J.** Representation of the analysis of cell polarity. Representative time lapse show that OS-stimulated macrophages maintain polarity when activated IN podosomes, while stimulation OUTside induces polarity changes (an effective polarization event is defined as a shift of 180° ± 40° between time points, bottom panel). **K.** Quantification of effective polarization events per cell reveals that OS repolarizes more when stimulated outside podosomes, whereas OH repolarizes when stimulated within podosomes (N=3; 40-50 cells/condition; statistical test: unpaired, Kuskal-Wallis with Tukey’s post hoc test). **L.** Analysis of podosome structure lifespan under local photo-stimulation shows that OH within podosomes leads to their disorganization, whereas OS disorganizes podosomes when stimulated outside of them (N=3; 40-50 cells/condition; mean ± SEM; statistical test: unpaired, Kuskal-Wallis with Tukey’s post hoc test). **M.** Schematic representation of subcellular photo-activation of OS and OH macrophages reveals how they regulate macrophage migration and cell polarization, and the synergy between both kinases to promote effective migration. Blue squares indicate photo-stimulation. All fluorescent photo-stimulation were performed at low stimulation frequency, 1 pulse every 240s (4mHz).Statistical significance: ns = no significant, *p<0.05, **p< 0.01, ***p< 0.001, ****p< 0.0001. Scale bar: 10 µm.

To test how the cooperation between podosome dynamics and clathrin-mediated endocytosis (CME) affects cell behavior, OS and OH activation was spatially controlled through digital micromirror device (DMD) photomanipulation; and distance over a fixed period of time, directionality and polarity (variation of angles between Front-Rear over time, Fig.5J) were quantified. In polarized RAW macrophages, activation of both optogenetic probes was controlled either in the podosome-rich front region (IN podosomes) or the clathrin hotspot-enriched rear/outer region (OUT podosomes, Fig. 5G). Spatial activation revealed strikingly opposing effects depending on both kinases and their subcellular localization.

When OS was activated in the front (IN), migration was enhanced, directionality remained high, and podosome structures were preserved, maintaining cell polarity (Fig. 5H-K, right graphs, movie S7). In contrast, activating OS at the rear (OUT) induced ectopic podosome formation, disrupting the existing macrophage polarization and, thus, impairing migration (Fig. 5H-K, left graphs, movie S8). These show that spatial confinement of SRC activity is key to sustaining directional migration. Conversely, OH activation showed the opposite pattern. When OH was activated in the front (IN), it disrupted podosome structures, repolarized macrophage, decreased directionality (decrease of MSD), leading to reduction of macrophage migration (Fig. 5H-K, right graphs, movie S9). On the other hand, activating OH at the rear (OUT) had no disruptive effect on podosomes, preserved macrophage polarity, promoting migration and directionality (Fig. 5H-K, left graphs, movie S10).

These findings support a model where macrophage migration relies on the coordinated, spatially segregated specific activities of SRC and HCK. At the front, SRC promotes podosome maintenance to sustain adhesion and protrusion. At the rear, HCK activation, through intense clathrin-mediated endocytosis sustained by specific activation of hotspots, induces a gradient disorganization of podosomes, causing them to form further away, which helps maintain podosome treadmilling and front-rear polarity (Fig. 5L). By being active at similar rates, OS and OH cooperate across distance, with the spatial separation of their specific functions enabling long-range functional integration. Rather than acting redundantly or in opposition, SRC and HCK function in a complementary and spatially integrated manner. Their distinct subcellular activities enable macrophages to balance migration and invasion in response to environmental cues, revealing an unexpected complexity in SFK signaling and its role in innate immune cell behavior.

### The adaptor modules of SFKs reveal the molecular basis of functional redundancy between SRC and HCK

Our optogenetic approach revealed that the specificity of SFKs signaling is primarily driven by the SH3-dependent nanoclustering. However, this raises the question of the origin of the molecular mechanism explaining overlapping functions of SRC and HCK *in vivo*.

To address this question, a modular design was used on optoSFKs to reconstruct step-by-step all the adaptor functionalities of each SFK: membrane anchoring, unique domain (SH4/UD) and SH2 domain. Indeed, our optogenetic SFKs are devoid of functional membrane anchoring and functional SH2 domains (Fig1 and ^27^).

Focusing on OH, which inhibits podosome formation under high-frequency stimulation (Fig. 2), the challenge was to determine how to reprogram its function to mimic the pro-podosome activity of SRC. Using the CRY2-CIBN system, which enables both homoligomerization and heteroligomerization, we systematically reintroduced modular features of the membrane anchoring and/or SH2 domains, fused to CIBN-GFP (Fig. 6A). Strikingly, only the combination of a palmitoylation-based membrane anchor (p59-palmytoylation or SH4-1 of HCK) with the SH2 domain of HCK was sufficient to restore OH’s ability to promote podosome formation, revealing a minimal modular requirement for functional redundancy. Neither module alone (SH2-CIBN or p59-CIBN) was sufficient. However, this functional redundancy of OH towards the SRC phenotype can be finely tuned. When OH was paired with the physiological myristoylation anchor of HCK (p61-myristoylation or SH4-2 of HCK), pro-podosome activity was not restored (Fig. 6B). This indicated that p59-HCK is the isoform responsible for functional redundancy with SRC, whereas p61-HCK contributes to distinct roles (confirming previous literature^11,42^.

**Figure 6:**
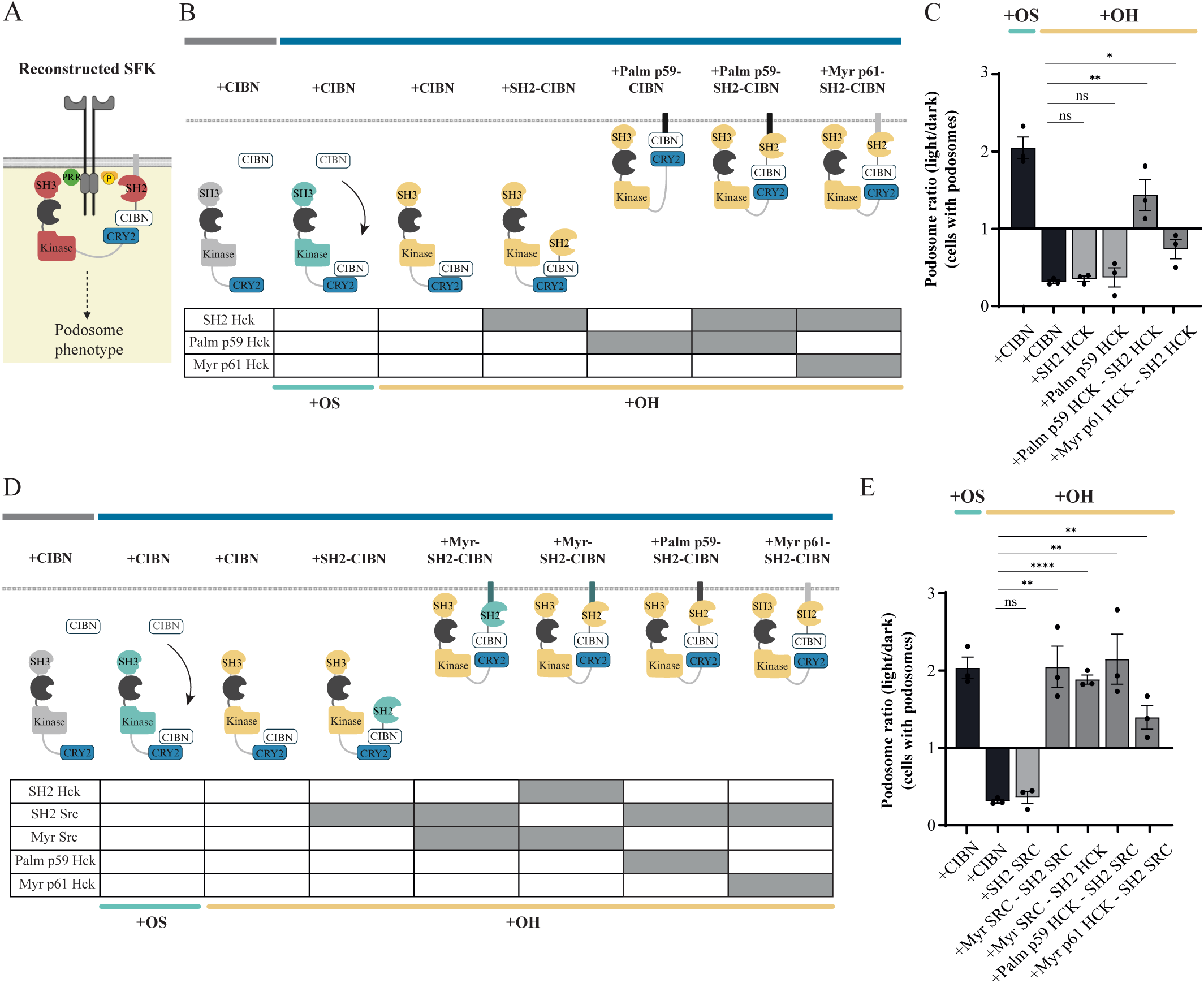
Hck chimeras reveal domain-specific redundancy with Src and conformation-dependent affinities for podosomes and clathrin structures. **A.** Schematic representation of a reconstructed SFK with a plasma membrane domain and SH2 linked to CIBN, which hetero-oligomerize with optoSFKs (SH3-kinase-Cry2) after photo-activation to form a complete chimeric SFK. **B-D.** Schematic representation of the different domain combinations chimeras (tagged with GFP) co-expressed with OS or OH in RAW 264.7 macrophages. **C-E.** Quantification of macrophages forming podosomes after 10 min of photo-activation with the indicated chimeras. The analysis reveals which domain combinations can modify the ability of activated OH to inhibit podosome formation. This analysis proposes to determine which domain combinations can bring back functional redundancy to activated OH, that is revealed by inducing an OS-like behavior such as podosome formation. Only the co-expression of SH4 and SH2 both fused to CIBN allows to induce a common function between OH and OS on podosome formation (N=3; 30 cells/replicate; data are presented as the ratio of cell with podosomes photo-activated vs. dark conditions; statistical test: unpaired, *t*-test test). All experiments of photo-stimulation were performed at high stimulation frequency, 1 pulse every 10s (100 mHz). Statistical significance: ns = no significant, *p<0.05, **p<0.005, ***p<0.0005, ****p<0.0001. Scale bar: 10 µm.

Interestingly, these SH2-membrane combinations also showed subtle specificity: SRC’s SH2 domain, when paired with myristoylation (SH4 of SRC), enhanced podosome formation more than HCK’s SH2 domain in the same context (Fig. 6B-C).

Altogether, the light-controllable modular domains approach demonstrates that functional redundancy between SRC and HCK is encoded in the combinatorial logic of two core features: membrane anchoring domain and SH2-mediated interaction. This approach opens new opportunities to dissect the biophysical and biochemical properties of SFK adaptor domains (membrane anchoring, UD-SH3 and SH2 domains) in a dynamic and tunable manner. The engineering of SFK domains allows to propose a new model of functional entanglement rather than simple redundancy or antagonism between SRC and HCK.

## Discussion

The molecular basis of signaling redundancy was revisited by focusing on the two Src Family Kinases (SFKs) that compensate the most for each other *in vivo*. Fully modular photoactivable SRC and HCK kinases were engineered to challenge the biochemical equilibria regulating their adaptor functions and kinase-dependent signaling activities. Our approach showed that SH3-dependent nanoclustering drives relocalization in specific sites of action for each kinase. In contrast, functional redundancy arises from the combinatorial integration of membrane anchoring and SH2 domains, which together restore shared signaling potential. This mechanism not only clarifies the basis of SFK redundancy but also reveals a previously unrecognized role for HCK in regulating polarized clathrin-mediated endocytosis through hotspots, impacting both directed migration and invasion in macrophages. Our findings support a model in which SRC and HCK are functionally entangled -exploiting both shared and distinct subcellular modules to cooperatively orchestrate complex cell behaviors.

### Seeking specificity between members of the SRC family

The balance between specificity and redundancy is a major question in cell signaling, particularly among kinase families. Redundancy is often invoked to explain why genetic loss-of-function approaches fail to yield distinct phenotypes across paralogous proteins or splice variants. However, while powerful, such methods are biased toward uncovering unique functions and often fail to capture the multifunctional and feedback-regulated nature of signaling proteins like SFKs. Among the ∼100 tyrosine kinases in the human genome^19^, SFKs stand out, comprising up to eight members in contrast to smaller families such as FAK or JAK with two to four members. Why evolution retained this level of duplication remains unclear. For 30 years, continuous efforts to define kinase specificity have focused on substrate profiling -via high-throughput phosphopeptide libraries^20,25^, analog-sensitive mutants^24^, and phosphoproteomics^43^- which have uncovered that the tyrosine kinases domain has some intrinsic preferences. Yet, due to their low catalytic activity, SFKs show limited substrate discrimination, making it challenging to derive functionally distinct roles from phosphorylation patterns^19–21^. Besides investigating kinase selectivity *in vivo*, specificity is thought to arise not from the kinase domain but from adaptor functions -principally through the SH2 and SH3 domains- which dictate subcellular localization and engagement with specific adaptor sets. Despite 50-70% sequence homology among SH2 and SH3 domains of SFKs^44^, prior reverse genetic studies have paradoxically shown that mutants of SRC deleted for either their SH2 or SH3 domain could rescue numerous phenotype^22,45–47^, suggesting their interchangeable properties in the adaptor functions. However, the importance of the SH3 domain could have been conclude evolutionary since the amino acid identity of the SH3 domain is more conserved than the SH2 domain in *Creolimax fragrantissima* that present the specificity to have a single non-receptor tyrosine kinase, CfrSRC^48^. Our optogenetic system presents the advantage of directly controlling the binding environment of the adaptor domains instead of simply controlling the opening of the kinase domain^29,30^, and revealed that specificity emerge from the light-induced nanoclustering of SH3 domains. Our observation reinforces the importance of the link between oligomerization of the SH3 domain and activation of SFKs as reported through the specific ability of the HIV-proteins NEF to activate HCK through its SH3 domain^49^, and the signaling properties of some SRC thermosensitive mutants dependent on their SH3 domain^46^. OS-induced SH3 nanoclustering was sufficient to localize SRC to focal adhesions or invadosomes -sites where proline-rich scaffolds concentrate- mirroring the behavior of endogenous SRC in both epithelial and mesenchymal cells (^27^ and Fig.2). This localization was not shared by HCK, despite its presumed redundancy. Instead, HCK nanoclustering targeted clathrin-coated structures (CCS), implicating the SH3 domain as a major determinant of spatial signaling specificity. Despite high homology between SH3 domains of SRC and HCK, this divergence could be further explained by the function of the unique domain (UD) of each SFK -an intrinsically disordered region (IDR) with low sequence homology across the family but high conservation across species for each member^23,50^. UDs have long been proposed to regulate substrate repertoires^51,52^, and our data now links them mechanistically to SH3 domain nanoclustering, reinforcing their role in defining context-specific interactions. By affecting the binding properties of the SRC SH3 domain^53^, the sequence specificity of each UD is thus directly connected with the SH3-dependent mechanism of specification of SRC. The link between IDRs and cooperative binding-such as phase transition mechanisms^54^- reinforces the importance of multivalent, cooperative interactions between SH3 domains with proline-rich proteins that could explain highly specific binding and selection of specific partners among PRR-containing proteins. Notably, this synergistic behavior is not observed with SH2 domains, underscoring the unique role of SH3 in defining subcellular localization and signaling outcomes. Together, it appears that specificity in SFKs is a modular, context-dependent property, shaped by SH3 nanoclustering, rather than by catalytic differences or static binding motifs alone.

### Optogenetic activation of HCK reveals a novel mechanism for regulating clathrin hotspots and polarized endocytosis function in macrophages

Beyond dissecting kinase specificity, our optogenetic approach uncovered an unexpected function of HCK in regulating a poorly characterized but specific subset of clathrin-coated structures known as *clathrin hotspots*^40^. Live-cell imaging combined with statistical analysis showed that they are characterized by repeated clathrin-mediated vesicle formation at the same membrane sites^55,56^, consistent with prior observations in neuronal cells^57^. Notably, the activation of OH induced localized highly curved clathrin assembly of 200 nm approximately distant from less 100nm to each other (Fig. 4), consistent with previous reports showing that controlled induction of highly curved membrane (200 nm radius) could induce repetitive assembly of clathrin and dynamin^58^. Electron microscopy further revealed that these assemblies are distinct from extended flat clathrin lattices^40^; instead, they appear as discrete, curved coated pits localized to specific membrane regions, possibly representing giant coated pits (GCPs) as described under conditions of elevated membrane tension induced by non-physiological osmotic shocks^59^. Importantly, HCK induces CCS with the same characteristic size as GCPs, highlighting a novel regulatory mechanism of endocytic hotspot formation. Moreover, the description of these clathrin hotspots reinforce the importance of spatially organized CME highly described in neurons^60^, and now essential to control macrophage functions. Indeed, inducing endocytosis at podosome-rich sites led to their disassembly, whereas activation at distal sites promoted maintenance of podosome polarity and oriented migration and front-rear polarity (Fig. 5). Hotspots may act as spatially confined endocytic hubs that avoid direct and total disruption of adhesive structures at the front of migrating macrophage and thus coordinate their turnover to induce a treadmilling process. This suggests that macrophages use spatially segregated trafficking to control adhesion structures and directional migration, as previously described in osteoclasts^61^.

Mechanistically, OH-clathrin hotspots are actin dynamic-dependent, as the pulsing of clathrin assembly was abolished by jasplakinolide treatment (Fig.5). While blocking fission with dynasore (dynamin inhibitor) did not prevent repeated clathrin assembly, but rather saturated hotspot capacity, highlighting a decoupling of initiation and vesicle scission at these sites. These findings suggest that specific optoSFKs such as OH can expose physiological regulatory mechanisms of CME that are otherwise obscured by the use of broad-spectrum inhibitors (e.g. PP2) or SFK mutants^62^. This functional link between HCK and clathrin hotspots is further supported by its known SH3 domain interactions with endocytic regulators such as dynamin^63^. Strikingly, OH failed to relocalize to clathrin structures in non-macrophage cell types (e.g. MDCKs, fibroblasts or HeLa), despite the presence of core endocytic components such as clathrin and dynamin2. This suggests that HCK does not target clathrin per se, but rather a distinct subpopulation of clathrin structures defined by a specific combinatorial set of associated proteins. Even within macrophages, OH labels only a subset of clathrin assemblies, providing another proof of the existence of molecularly distinct clathrin subpopulations whose composition and function remain unresolved. These observations clearly highlight that clathrin-coated structures are not uniform across cell types -or even within the same cell (Fig.3–4), as previously observed with the differential endocytosis of splice variants of the GPCR able to bind actin^64^. Thus, our optogenetic HCK approach offers a novel tool to dissect the molecular determinants and functional specialization of these heterogeneous endocytic domains.

### Redundancy or functional entanglement

This nanocluster- and SH3-dependent mechanism of specification of SFKs sustains a conceptual framework where highly homologous kinases can have distinct biological roles while retaining potential for functional compensation under specific conditions such as those induced by genetic loss-of-functions experiments. The link between nanoclustering and activation is a general mechanism observed for numerous signaling elements active such as Ras or Rac GTPases^65,66^, and even SFKs^17,18^. By controlling nanoclustering and uncoupling SH3 and SH2 binding of SFKs, our optogenetics does not create artificial properties of SFKs but allow to uncouple specific and common functions of SFKs. However, the comparison between optoSFKs and endogenous SFKs raised the question of the equilibrium between both type of functions for SFKs. Indeed, macrophages can express up to 6 members (SRC, FYN, YES, HCK, FGR and LYN) that co-exist and appear being able to be localized either in the same subcellular structures and in specific ones. For example, a single member as HCK will be both in adhesion sites and in clathrin structures. This coexistence can now explain the observed discrepancy of HCK subcellular relocalization reported by different laboratories^11,67^. Our multimodular reconstruction of the adaptor functions of SFKs (membrane anchoring-SH3-SH2) thus proposes that the biochemical equilibrium between each domain should drive the probability to be accumulated in specific subcellular structures or not. This implies that within a single SFK population, distinct subpopulations can emerge, enabling co-accumulation with other SFKs in shared compartments (e.g. HCK with SRC in adhesions), or distinct localization separate zones (e.g. HCK in clathrin hotspots). The existence of both specific and common binding partners mediated by two different pools of the same molecules support that the notion of redundancy is not explained by simple functional overlap. On the contrary, the absence of a phenotype observed downstream loss-of-function genetic experiments is not due to a simple redundancy mechanism but could be nicely explained by a functional entanglement of different SFK members presents both in common sites of action but also specific sites. One advantage of having the same family of signaling regulators in distinct and separated subcellular structures is that their signaling activities are naturally in phase since having the same range of kinetic parameters of activation-inhibition in each subcellular structure. As a result, signaling outputs become more robust and can lead to cooperation between different subcellular structures, even at long distance, in order to develop complex cellular processes such as migration and cell invasion. The description of this functional entanglement mechanism invites revisiting the description of unexplained cooperative signaling pathways that can act more efficiently only when altogether.

In conclusion, exploration of the redundancy between SFKs revealed new insights into fundamental aspects of cell signaling and paves the way to develop a conceptual framework for new biotechnology approaches controlling specific immunological functions of macrophages.

## Movie legends

Movie Sup.1 : Live TIRF imaging (legend integrated in the movie + time stamper) of macrophage where OS or OH are photoactivated (100mHz imaging frequency and activation).

Movie Sup.2 : Live TIRF imaging (legend integrated in the movie + time stamper) of MDCKs where OS or OH are photoactivated (50mHz imaging frequency and activation).

Movie Sup.3 Live TIRF imaging (legend integrated in the movie + time stamper) of macrophage expressing LifeAct-GFP where OS is photoactivated (500mHz imaging frequency and activation).

Movie Sup.4 Live TIRF imaging (legend integrated in the movie + time stamper) of macrophage expressing LifeAct-GFP where OH is photoactivated (500mHz imaging frequency and activation).

Movie Sup 5 : Live TIRF imaging (legend integrated in the movie + time stamper) of macrophage expressing CLC-GFP where OH is photoactivated (100mHz imaging frequency and activation).

Movie Sup 6 : Live TIRF imaging (legend integrated in the movie + time stamper) of macrophage expressing CLC-GFP where OS is photoactivated (100mHz imaging frequency and activation)

Movie Sup 7 : Live EPIfluorescence imaging (legend integrated in the movie + time stamper) of macrophage migrating when OS is activated only in its front (IN podosomes-blue segmentation, 4mHz imaging frequency and activation)

Movie Sup 8 : Live EPIfluorescence imaging (legend integrated in the movie + time stamper) of macrophage migrating when OS is activated only in its rear (OUT podosome-blue segmentation, 4mHz imaging frequency and activation).

Movie Sup 9 : Live EPIfluorescence imaging (legend integrated in the movie + time stamper) of macrophage migrating when OH is activated only in its front (IN podosome-blue segmentation, 4mHz imaging frequency and activation)

Movie Sup 10 : Live EPIfluorescence imaging (legend integrated in the movie + time stamper) of macrophages migrating when OH is activated only in their rear (OUT podosome-blue segmentation, 4mHz imaging frequency and activation)

## Material and Methods

### Experimental cell models and culture

Murine RAW 264.7 macrophages cell line was transiently and/or stably transduced using a viral and/or transposon strategy. Cells were cultivated at 37⁰C and 5% CO_2_ in RPMI 1640 medium supplemented with GlutaMAX™ (Gibco, Thermo Fisher Scientific), 10% fetal bovine serum (FBS) and 1% (v/v) penicillin-streptomycin (P/S). Transient transfection of RAW 264.7 macrophages was performed using jetPRIME® transfection reagent (Polyplus-transfection) with optimized conditions (0.8 µg plasmid DNA, 75 µL jetPRIME® buffer, and 1.6 µL reagent, in a final volume of 700 µL).

Primary mouse Bone Marrow-Derived Macrophages (BMDM), primary human macrophages, epithelial MDCK cells, epithelial human breast cancer MDA-MB 231 cells, MEF fibroblasts, epithelial human cervical cancer HeLa cells, and SYF fibroblasts (Src-/-, Yes-/-, and Fyn-/-) were used. BMDM cells were isolated from mice bone-marrow, primary human macrophages were derived from patient blood samples (EFS Grenoble) and cultivated at 37⁰C and 5% CO_2_ in α-MEM medium (Gibco, Thermo Fisher Scientific) supplemented with 10% FBS, 1% P/S, and 20 ng/mL Macrophage colony-stimulating factor (M-CSF, R&D systems). M-CSF was replaced each 2 days with complete media. BMDM and human macrophages were electroporated (Invitrogen™ Neon™ Transfection System; MPK5000) with 1 µg plasmid DNA, 10 µL reagent R, and 1000 V_40_2 pulses.

The rest of the cells were cultivated at the same conditions in DMEM, high glucose, GlutaMAX™ supplement (Gibco, Thermo Fisher Scientific) with 10% FBS and 1% P/S. These cells were transfected using either Lipofectamine 2000 according to the manufacturer’s protocol (Invitrogen) or jetPRIME.

### DNA constructs and plasmids

In this study, the various Optogenetic SFKs plasmids are the result of C-terminal fusion of CRY2–mCherry to the indicated SFK mutants. These, and additional recombinant DNA constructs are listed in key resource table.

Plasmids were constructed using Gibson Assembly, according to the manufacturer’s instructions (New England Biolabs, NEB), made nicely available by the indicated principal investigator through purchased from Addgene (see details articles and addgene number in key resource table) or kindly given as gifts by the indicated collaborators.

### Obtention of stabling expressing RAW 264.7 macrophages

#### Lentiviral infection

Lentiviruses were produced by co-transfecting pC57GPBEB GagPol MLV, pSUSVSVG (gifts of Dr Negre, ANIRA Platform, SFR Biosciences, Lyon, France) and each plasmid of interest using Lipofectamine2000 (Invitrogen) in HEK293 FT cells (gift of Dr Negre) plated in 6-well plates at 50%’ confluency. The medium was changed 24 h later. The viral supernatant was collected after 72 h and was filtered with 0.45 μm filters.

RAW 264.7 cells were plated in 6-well plates so as to achieve 60% confluency on the day of infection. The filtered supernatant was directly used to infect cells of interest. The medium was changed 24 h after infection. After 10 days of decontamination, cells were FACS sorted (Aria cell sorter 2000, BD) based on the level of expression of mCherry-tagged OSFK in order to have no dark binding of the CRY2 system, using excitation with a 561 nm LASER.

#### Transposon

GFP-tagged endocytic markers were stably expressed in RAW 264.7 cell lines stably expressing optoSFKs using the Sleeping Beauty transposon system. Briefly, cells were seeded one day prior to transfection, at approximately 50% confluency. Transfection was performed following the previously mentioned jetPRIME protocol, in antibiotic-free RPMI medium. Plasmid DNA encoding the marker of interest was co-transfected with the Sleeping Beauty transposase plasmid at a 1:10 ratio (total DNA). After 5–6 hours, the transfection medium was replaced with complete RPMI medium supplemented with 10% FBS and 1% P/S. Puromycin selection (5 µg/mL) was initiated 48 hours post-transfection and maintained by replacing the medium every 2 days until all non-resistant cells were eliminated.

### Single cells optogenetics experiments and TIRF imaging

Live-cell imaging was performed on an inverted motorized microscope (Zeiss AxioVert 200M) equipped with a Coolsnap HQ2 camera (Photometrics), 100× (NA 1.46, oil) Plan-Apochromat oil objective (Rapp Optoelectronic). TIRF illumination was performed using DPSS lasers at 488 nm, and 561 nm, combined with a Slider TIRF2 module (ZEISS). During imaging, cells were placed on a heated 37°C stage (ZEISS) combined with an incubator of CO_2_ (XL incubator, PeCon). Acquisition and experimental control were managed with MetaMorph software (Universal Imaging).

Image brightness and contrast were adjusted post-acquisition using ImageJ to enhance visualization of optogenetic probe recruitment. For photo-stimulation experiments, brightness settings were standardized across conditions and optimized based on the maximal intensity observed in the stimulated images. This adjustment was applied uniformly to allow accurate comparison and to highlight probe dynamics upon activation.

### Optogenetics experiments on large population activation

#### 2D migration

For optogenetics stimulation on large cell population in 2D migration assays, cells were seeded one day prior to imaging and starved in serum-free medium for at least 1 hour before the experiment. Live-cell imaging was performed using the same inverted microscope described above, equipped with a 20×/0.5 Achromat objective. Transmission images were acquired every 4 minutes for 3–4 hours, using a Colibri lamp and epifluorescence filters. Red channel images captured in parallel to identify OptoSFK-positive cells. Subsequently, blue-light illumination was applied to photoactivate the cells, and time-lapse imaging was continued for up to 16 hours.

#### 3D invasion - xCELLigence

Three-dimensional invasion assays were performed using the xCELLigence RTCA DP Real-Time Cell Analyzer – Dual Purpose (Agilent) in combination with 16-well CIM-Plates (ACEA, Agilent). Matrigel (reduced phenol red, Corning, Cat# 356231) was diluted to 3.3% in serum-free RPMI and added to the upper chamber of the CIM-Plates. Plates were incubated overnight at 37 °C to allow Matrigel polymerization. Cells were then seeded one day prior to the experiment and starved in serum-free medium for at least 6 hours before the assay. On the following day, RPMI supplemented with 10% fetal bovine serum (FBS) was added to the lower chamber as a chemoattractant. Cells were seeded in the upper chamber in serum-free RPMI. Real-time impedance-based measurements of cell invasion were acquired every 4 minutes over a 36-hour period. For photoactivation conditions, cells were illuminated every 4 minutes (4 mHz) using the LITOS LED device and disposed on the top of the CIM plates.

### Subcellular photoactivation

Live-cell imaging was performed on an inverted motorized microscope (Nikon Ti-E) equipped with a Zyla camera (Andor) and a 60× Plan-Apochromat oil objective (NA 1.4, Nikon). Illumination was provided by SpectraX LEDs (Lumencor) at 475/25 nm and 645/30 nm. For light patterning, a Mosaic 3 digital micromirror device (Andor) was used. Cells were maintained on a heated 37 °C stage (Nikon) with CO₂ control via an environmental chamber (Life Imaging Services). Image acquisition and experimental control were managed using custom scripts built on Pycro-Manager (http://dx.doi.org/10.1038/s41592-021-01087-6) and μManager (http://dx.doi.org/10.14440/jbm.2014.36), as described previously (https://doi.org/10.7554/eLife.90305.1). Cell segmentation was performed using a Cellpose 3 model finetuned on custom data (http://dx.doi.org/10.1038/s41592-025-02595-5), and podosome detection was carried out with a custom-trained Mask2Former instance segmentation model. Cell tracking was conducted using the TrackPy library (https://zenodo.org/doi/10.5281/zenodo.12708864). For IN podosome stimulation, masks were generated directly from the podosome segmentation. For OUT podosome masks, the cell’s major and minor axes were first determined. The front of the cell was defined as the side of the major axis where a podosome region was detected. The stimulation mask was then created as a line parallel to the minor axis, positioned at a fixed percentage behind the front along the major axis.

Podosome segmentation model available online (CC BY 4.0 license):

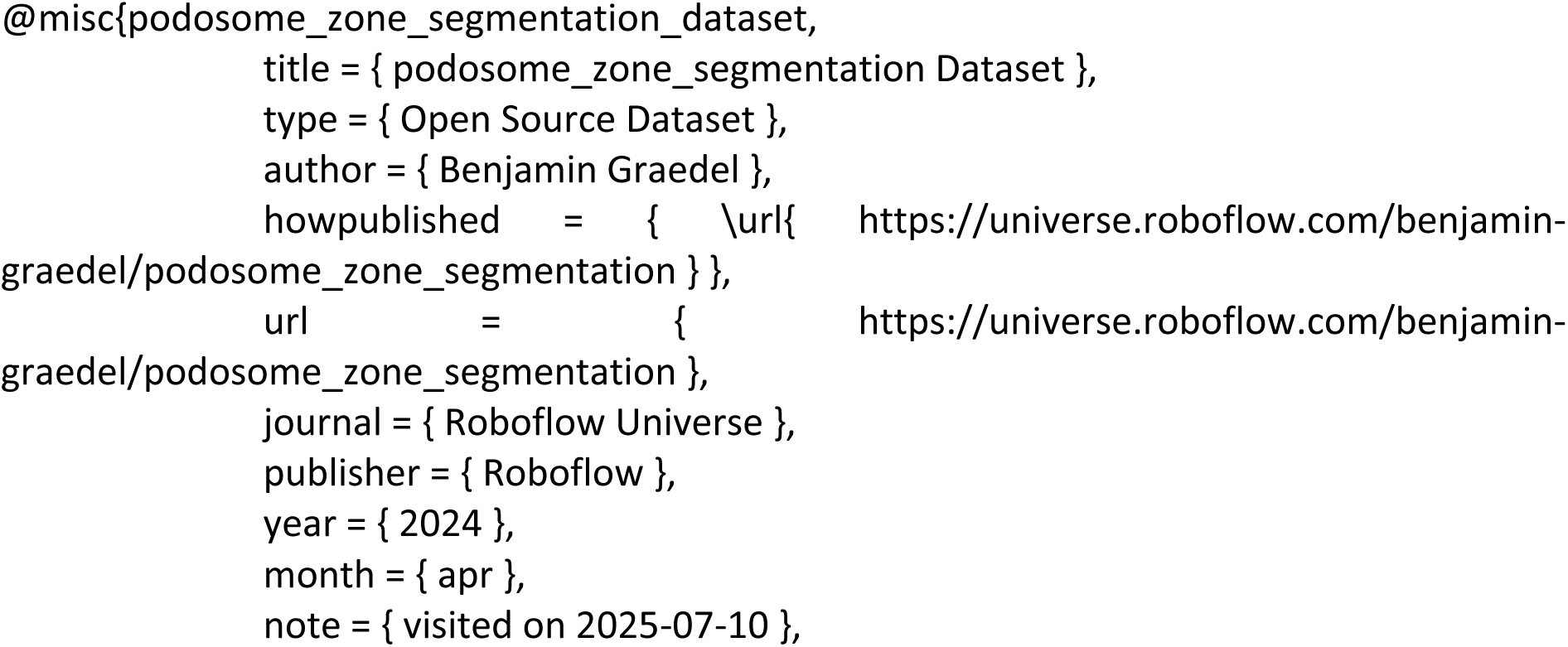

### Western blotting

Protein phosphorylation levels were assessed by Western blot analysis. For control conditions involving Src family kinase (SFK) inhibition, cells were treated with 20 μM PP2 (Calbiochem, Cat# 529576) for 30 minutes prior to photo-stimulation. Light stimulation of large cell populations was performed using a LITOS system LED plate from Pertz lab in Bern, Switzerland (T. Höhener et al. 2022), delivering blue light pulses every 10 seconds for 15 minutes before cell lysis. Cells were lysed in a modified RIPA buffer [150 mM NaCl, 20 mM Tris-HCl (pH 7.4), 1% NP-40, 0.5% sodium deoxycholate, 0.1% SDS, 1× EDTA-free protease inhibitor cocktail (Sigma, Cat# 11836170001), and a phosphatase inhibitor mix containing 1 mM sodium orthovanadate, 10 mM sodium fluoride, and 10 mM β-glycerophosphate]. Lysates were subjected to immunoblotting using the following primary antibodies: STAT3 (clone 124H6, Cell Signaling Technology, Cat# 9139), phospho-STAT3 (Tyr705, clone D3A7, Cat# 9145), phosphotyrosine antibodies 4G10 (Millipore, Cat# 05-321) and PY100 (Cell Signaling Technology, Cat# 8954), and GAPDH (Cell Signaling Technology, Cat# 2128). All primary antibodies were used in 5% BSA in TBS-Tween (TBS-T). Western blot images were acquired using a Vilber Newton 7.0 system, and densitometric analysis was performed using ImageJ software with the Gel Analysis plugin. Band intensities were quantified and normalized to GAPDH to determine relative protein expression levels.

### PREM and CLEM Electron Microscopy

Adherent plasma membranes of murine RAW 264.7 macrophage cell lines on glass coverslips were disrupted by sonication according to Heuser (2000b). The unroofed cells were then fixed and processed as previously described (Vassilopoulos et al. 2019). Sequential treatments involved 0.55% OsO4, 1% tannic acid, and 1% uranyl acetate, followed by graded ethanol dehydration and substitution with hexamethyldisilane (Sigma). Once dried, the samples were rotary shadowed with 2 nm of platinum and 6 nm of carbon using a high-vacuum sputter coater (Leica Microsystems). The resulting platinum replica was removed from the glass with 5% hydrofluoric acid, thoroughly washed in distilled water, and mounted on 200 mesh formvar/carbon-coated EM grids. The images were captured with a digital camera (Xarosa) attached to a transmission electron microscope (Joel, USA) operating at 120 kV, with the grids mounted on an eccentric side-entry goniometer stage. The images were adjusted for brightness and contrast using Adobe Photoshop (Adobe, USA) and presented in inverted contrast. For CLEM, unroofed and stained samples were imaged by spinning disk microscope, prior to make the replica. Grids were first imaged by low-magnification EM to relocate the light microscopy-imaged area, followed by high-magnification EM of the corresponding ROIs, and aligned using affine transformation via the eC-CLEM plugin in ICY^68^.

### Morphometric analysis of CCPs by PREM

To study structural details of CCPs we performed a morphometric analysis in which clathrin structures present on PREM images were plotted according to their area and curvature after measuring each structure’s electron opacity in OS and OH without and with 10 minutes of photo-activation as described previously^69^. We defined an area threshold cutoff corresponding to the size of the largest CCPs encountered with diameters of ∼ 8 µm^2^, while a curvature threshold was set manually to segregate the clathrin-coated pits respectively.

To calculate the area of ROIs, each clathrin structure is manually traced based on its honeycomb appearance. The mean gray level of each ROI is quantified as a substitute for structure curvature. The spherical pits on the y-curvature axis correspond to lighter gray hues. Each clathrin structure is plotted on an x-y graph, with area on the x-axis and curvature on the y-axis. Two cutoff thresholds are employed to clearly distinguish small and large CCPs. Figures 4 and supporting figure 4 are related.

### Transferrin uptake assay

Cells were seeded the day prior to the experiment in ibidi 4-well µ-slides and starved in serum-free RMPI for 30 minutes before the experiment. Alexa Fluor 488-conjugated transferrin (Invitrogen, Cat# T13342) was prepared at a concentration of 50 µg/mL in FluoroBrite DMEM (Gibco), a phenol red-free medium optimized for fluorescence imaging. Cells were incubated with transferrin on ice for 5 minutes, then incubated with for 0min or then placed at 37°C with 5% CO_2_ for 15 minutes, with or without photoactivation (blue light stimulation every 10 seconds, using LITOS system), washed once with 1% BSA/PBSA and fixed with 4% paraformaldehyde (PFA). Following incubation, cells were washed and fixed under the same conditions as the 4°C control. Imaging was performed using TIRF microscopy, and 30 cells were imaged per condition. For quantification, mean fluorescence intensity per cell was measured in ImageJ and normalized to background fluorescence.

### Image analysis

#### Co-occurrence analysis

Co-occurrence analysis was performed using the ComDet plugin(Eugene Katrukha / https://github.com/UU-cellbiology/ComDet/wiki / in ImageJ as described in Fig S1 C. The plugin was configured to detect spots based on a defined particle size and intensity threshold. Colocalization events were identified based on spatial proximity between detected puncta in the two channels within a given maximal distance (3-4 pixels). The percentage of co-occurring spots was calculated relative to the total number of spots in each channel.

#### Quantification of OptoSFK probe relocalization

To assess the recruitment of the OptoSFK probe, normalized intensity was performed on individual cells using ImageJ. For each cell, the minimum and maximum fluorescence intensities were measured. The maximum intensity - corresponding to the recruitment site – was divided by the minimum intensity, which was used as an estimate of the background signal. This normalization method was selected due to the high background levels that made average intensity measurements unreliable and unable to reflect localized recruitment. By comparing the max-to-min intensity ratio, recruitment efficiency could be quantified in a background-independent manner.

#### Quantification of Podosome and focal adhesion

Podosome and focal adhesion analyses were performed using ImageJ. Podosomes were detected using the TrackMate plugin (https://imagej.net/plugins/trackmate/; Ershov, D., Phan, M.-S., Pylvänäinen, J. W., Rigaud, S. U., Le Blanc, L., Charles-Orszag, A., … Tinevez, J.-Y. (2022). TrackMate 7: integrating state-of-the-art segmentation algorithms into tracking pipelines. *Nature Methods*, *19*(7), 829–832. doi:10.1038/s41592-022-01507-1) by applying an intensity threshold to identify individual structures. The number of podosomes per cell was normalized to the initial time point (time 0) to assess dynamic changes over time. Podosome lifespan was measured by tracking individual structures using TrackMate, and the average track duration per cell was calculated and plotted. Classification of podosome organization (single vs. ring) was performed manually upon photo-stimulation: single podosomes were defined as individual F-actin puncta, whereas ring structures were characterized by a circular arrangement of multiple individual podosomes.

Focal adhesions were manually quantified using similar thresholding, with counts also normalized to time 0. Focal adhesion length was measured manually by drawing a line along each adhesion and recording its length using ImageJ’s measurement tools.

#### Spot and cell migration tracking

Spot tracking was performed using the TrackMate plugin in ImageJ. Spots were detected based on intensity thresholding and particle size settings, which were kept consistent across all conditions. For each detected spot, the lifespan and maximum displacement were calculated and visualized using Kernel Density Estimation (KDE) plots. Short tracks comprising fewer than five frames were excluded from the analysis to eliminate noise and ensure reliability.

- For macrophage migration analysis, manual tracking was performed using the MTrackJ plugin (https://imagescience.org/meijering/software/mtrackj/; E. Meijering, O. Dzyubachyk, I. Smal <u>Methods for Cell and Particle Tracking</u> *Methods in Enzymology*, vol. 504, February 2012, pp. 183-200) in ImageJ, enabling frame-by-frame tracking of cell trajectories. Mean square displacement (MSD) was calculated using the Excel macro provided in the study of (Gorelik et al. 2014).

#### Endocytic frequency

To quantify endocytic frequency, kymograph analysis was performed from time-lapse videos using ImageJ. Kymographs were generated by drawing a line across regions containing OptoSFK and/or clathrin-GFP spots. Line selections were made to follow the path of dynamic structures over time. For each line, a plot profile was extracted, where the x-axis represents time, and the y-axis corresponds to fluorescence intensity of both optoSFK and clathrin-GFP. This allowed visualization and quantification of the temporal dynamics of individual endocytic events. The frequency of events was assessed based on the number of intensity peaks over time, representing successive cycles of clathrin-mediated endocytosis.

The analysis was performed with a custom Python script relying mainly on the PeakUtils library (https://peakutils.readthedocs.io/en/latest/). Thus, the traces were corrected by baseline detection and subtraction. Traces were further smoothed by Lowes filtering. Peak detection was performed with the same settings for each trace and for all conditions. Peak frequency and inter-peak distances were extracted from this data.

#### Effective polarization analysis

Effective polarization analysis was performed manually using ImageJ. A line was drawn from the center of the cell to the leading edge to define the polarization axis, and the angle of this axis was measured using Radius measurements. The change in polarization direction between consecutive frames was calculated by determining the difference in angles. An effective polarization event was defined when the angular shift between frames was approximately 180° (±40°), accounting for variability due to manual measurement (Fig.5J). This approach allowed the identification of significant polarity reversals during cell migration.

#### Statistical analysis

Statistical analyses were performed using GraphPad Prism 10 (GraphPad Software). Statistical significance was defined as P < 0.05. Specific statistical tests used are indicated in the figure legends. Data are presented as mean ± standard deviation (SD) or standard error of the mean (SEM), as specified. Significance levels are denoted as follows: ns, not significant (P > 0.05); P < 0.05; P < 0.01; *P < 0.001; P < 0.0001.

Schematic figures were created using BioRender. Statistical analyses and graph generation were performed using GraphPad Prism (version 10) or Seaborn. Final figure assembly and image formatting were carried out in Adobe Illustrator. The main text was written by CTT and OD, then submit to Mistral AI and ChatGPT for wording optimization.

## Acknowledgements

This work was funded by ANR “NODES” and ARC programs (coordinator: O.D.), by LLNC for PhD funding of C.T.T., P.R. and L.C. and by AC. 4^th^ of PhD of C.T.-T. was funded by ARC. We thank the strong and efficient support of the imaging platform MicroCell.

## Authors Contributions

C.T.-T., P. R.., L.C., C.O. and O.D. generated constructs and cell lines, performed and analyzed data in cell biology. C.T.-T. and F.S. developed co-occurrence, spots density plots and peak analysis. B.G. and L.H. developed spatially controlled activation in segmented region of interests (IN or OUT podosomes). A.G. support the excellence of TIRF imaging on the MicroCell platform.

C.T.-T., P. R. and O.D designed the study, and C.T.-T., and O.D. wrote the paper with significant contributions from all authors.

